# High-throughput bacterial aggregation analysis in droplets

**DOI:** 10.1101/2024.09.24.613170

**Authors:** Merili Saar-Abroi, Karoliine Lindpere, Daniel Kascor, Triini Olman, David Gonzalez, Fenella Lucia Sulp, Katri Kiir, Immanuel Sanka, Simona Bartkova, Ott Scheler

## Abstract

Microfluidic droplet platforms provide a rapid tool to study and capture bacterial aggregation in a well-controlled micro-environment, while image analysis presents an easily available technique to investigate droplet contents. However, the lack of standardised, well-documented methods and reliance on custom image analysis workflows limits wider adoption of the method and produces inconsistent, incomparable data on aggregation. We present a robust, cost-effective method using both mono- and polydisperse droplets and texture-based image analysis via an open-source software CellProfiler™ to assess bacterial aggregation. Compared to a manual droplet evaluation carried out by a human expert panel, textural characterisation achieves accuracy over 90% and more than 80% precision. Applying the pipeline, we found that suboptimal antibiotic concentrations can increase aggregation, whereas exposure to microplastic beads and metals reduces it. Overall, the developed pipeline offers high accuracy, easy setup, and broad applicability for bacterial aggregation.

## Introduction

Antimicrobial resistance (AMR) is a global threat that can result in millions of deaths if preventive measures are not implemented. The global prevalence of deaths associated with bacterial AMR rose to 4.71 million in 2021, and a progressive upsurge is projected to reach up to 8.22 million deaths due to AMR in 2050 [1].

Bacterial aggregation and biofilm formation is seen as one of the key concerns in addressing AMR [2]. Aggregated lifestyle of bacteria ensures the survival of the innermost bacteria in this multicellular matrix even in hostile environments, while simultaneously providing a fruitful ground for horizontal gene transfer [3]. Once unfavourable conditions subside, bacteria could separate from the aggregate matrix and successfully colonise new surfaces [3]. This phenomenon is known to be fueled by constant antibiotic exposure and the presence of contaminants as aggregation platforms [4]-[5].

Altogether, these figures underscore a critical need for the exploration of novel techniques to monitor and study bacterial aggregation and AMR evolution. As the burden of AMR is considered the greatest in low-income and middleincome countries, cost-efficient and innovative techniques pleasing a variety of surveillance strategies of antimicrobial resistance are in high demand [6].

The most common methods for bacterial aggregation studies are sedimentation assays or flow cytometry [7][8]. However, these methods have the major limitation to be time-consuming or expensive [9]. Water-in-oil droplets, acting as nanoscale bioreactors, exhibit great potential at increasing the precision of microbiological experiments [10]. Furthermore, the method is much faster and enables identifying bacterial growth within 3–4 hours as opposed to 16–20 hours required by the traditional culturing methods [11].

Recent advancements have employed droplets for a wide range of microbiological assays and experiments. The most promising use of droplets is for antibiotic susceptibility testing, which would enable rapid evaluation of bacterial responses to antimicrobials [12]. Droplet platforms are also advantageous in enzyme activity screening, metabolic profiling and microbial interaction studies, where confined microenvironment allows for precise reaction conditions and bacterial behaviour observation [13]. Lastly, the platform exhibits great potential to isolate and cultivate rare or slow-growing bacteria, that thrive only in delicately configured environmental conditions [13].

Two main methods of creating water-in-oil droplets are recognised, and each approach comes with trade-offs in ease of use, reproducibility, and analysis.

Using monodisperse droplets for biological experiments offers high reproducibility between experiments and precise control over droplet size and volume, leading to easier detection and analysis workflows [14], [15], [16]. However, producing them requires advanced laboratory equipment and the purchase of a commercially available microfluidic droplet system may introduce financial and training-related challenges for the user interested in the technology [17].

As another option, droplets could be generated in a polydisperse distribution through a simple vortex of immiscible and aqueous liquids [17]. A significant drawback of polydisperse droplets is seen most in lack of precise volume control leading to batch-to-batch variability, higher surfactant consumption, and more complex software algorithms to accurately detect and analyse droplets [15], [16], [18]. Despite abovementioned limitations, polydisperse droplets provide the same throughput and droplet stability over time compared to monodisperse droplets, making it a great, highly accessible alternative to monodisperse droplet systems [17].

Droplet experiments have been coupled with many diverse modalities of droplet detection and analysis, optical detection being the most common approach. Depending on experiment nature, one could analyse droplets using microscopy imaging, Raman spectroscopy, and many more [19]. Regarding aggregation, imaging provides direct insight in the dynamics of such bacterial behaviour. Furthermore, imaging equipment is commonly available in laboratories, making droplet analysis highly accessible for anyone.

Due to high-throughput nature of droplet-based techniques, visual inspection of images is laborious and therefore, not a viable option for droplet image analysis. So far, automated image analysis often relies on high proficiency in programming skills using Python, MATLAB, and other advanced tools [20]. These proficiencies may fall outside the typical training of many biomedical professionals, preventing their widespread adoption in droplet analysis.

Alternatively, commercially available sophisticated software for image analysis may aid in this issue. However, the use of these more sophisticated image analysis tools is often hindered by high cost, leaving researchers little options to set up a droplet analysis workflow without significant financial investment.

Lastly, a variety of script-free open-source software have been developed to assess different biological structures, such as CellProfiler™, ImageJ, ilastik, QuPath, and many more. The uniform morphology of droplets makes it easy to adapt such software for droplet detection and consecutive droplet content analysis. Previously, Sanka et al. (2021) studied the suitability of such software for droplet detection and found high accuracy, precision and wide applicability of CellProfiler™ for various droplet experiments compared to other similar software [20].

Although such script-free open-source software demand little in terms of finances, training and technical setup, the lack of readily available workflows with thorough documentation tailored specifically for droplet analysis remains a significant obstacle in mainstream adoption |droplet technologies

Therefore, a clear need to bridge the gap between advanced bioinformatics and droplet image analysis for broader user base comes sharply into focus. In this paper we propose a solution to this need by using water-in-oil droplets in combination with user-friendly image analysis tools to assess the aggregation patterns of bacteria in high-throughput. The applicability of the method is demonstrated by easy-to-use image analysis pipelines established using a free, open-source software CellProfiler™ to quantify bacterial aggregation in response to suboptimal concentrations of antibiotics, microplastic beads or metals in both mono- and polydisperse droplets. The pipelines presented in this paper are accompanied with thorough guidelines including troubleshooting instructions for easy adaption of the method. Lastly, we seek to encourage the discussion and the development of more user-friendly and transparent analytical tools for droplet research.

## Results and Discussion

An image analysis pipeline utilizing textural characterisation within CellProfiler^™^ was specifically designed to identify and aid classifying droplets based on bacterial state (‘no growth’, homogenous growth’, ‘aggregation’) detected inside droplet (Figure 1a). Texture is a feature describing the variability and spatial arrangement of pixel intensities across an image or an object. In single-cell state of life, bacteria are evenly distributed in droplets, resulting in little difference between pixel intensities across droplet. In aggregation, pixel intensities vary greatly in droplets - pixels in areas emptied due to bacterial migration or death show low or no signal, while aggregated regions exhibit strong intensity in proximity.

**Fig. 1.**
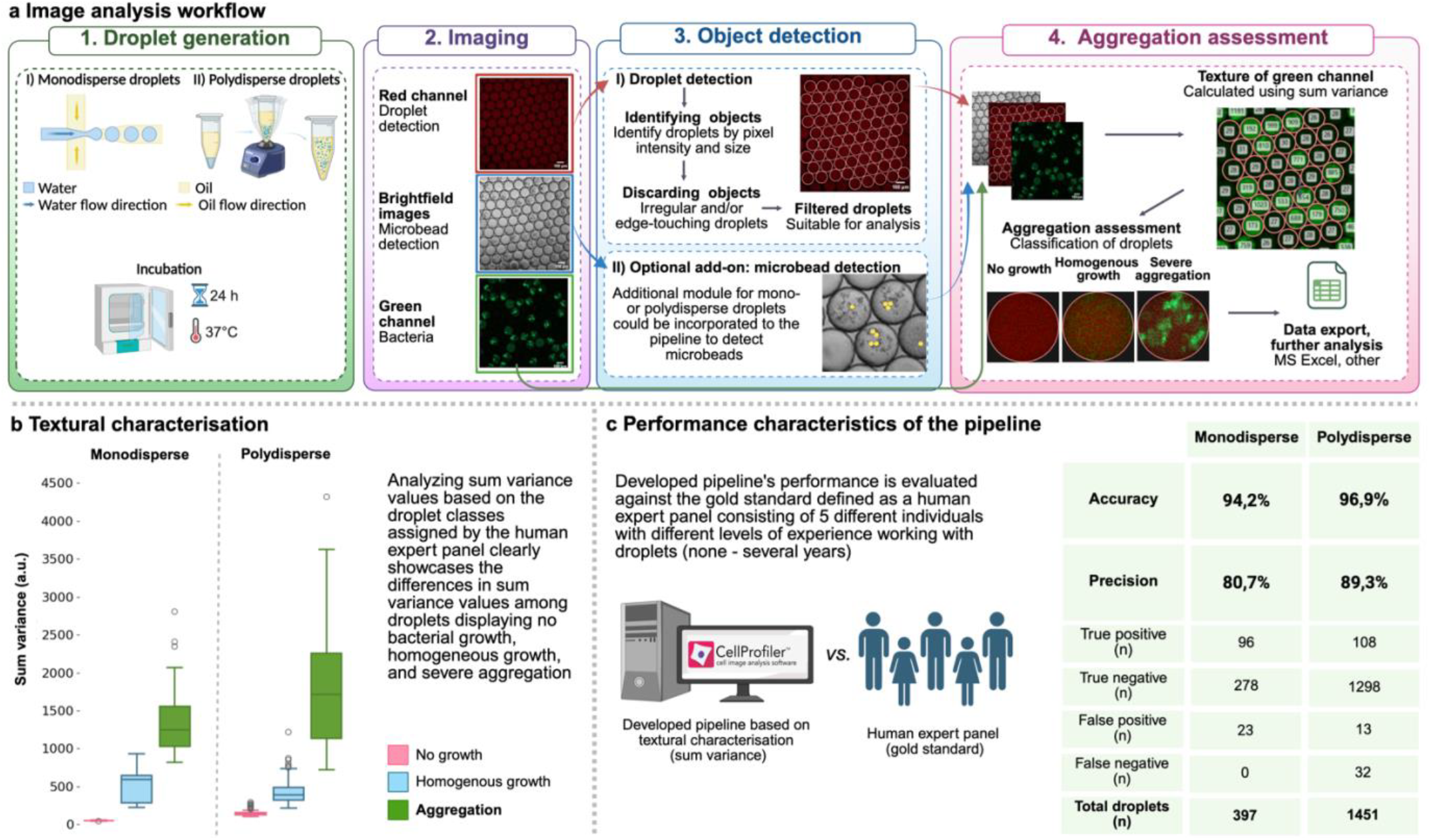
Droplet generation and image analysis workflow utilising textural properties in CellProfiler^™^ showcases a fit tool for bacterial aggregation evaluation in droplets. **a** Protocol overview of the bacterial aggregation assessment using CellProfiler™. Droplets of mono- or polydisperse nature can be used to analyse bacterial aggregation. Generation of droplets is followed by further incubation to encourage bacterial growth and study the phenomenon under investigation. Droplets are visualised via microscopy, where three types of images are acquired – brightfield, red, and green channel. Droplets are detected using the red channel of the images, microbeads via brightfield images and textural characterisation is performed in the green channel. All data gathered during the analysis is exported as a .csv file and could be analysed using any spreadsheet software. **b** Higher values of textural characteristic - sum variance - measurements reveal severely aggregated droplets. Shown data is applicable to experiments performed with sub-optimal concentrations of cefotaxime in mono- and polydisperse droplets. Boxplots display the minimum, first quartile, median, third quartile and maximum values of sum variance in the classification group. Outliers above interquartile range of 1.5 are shown as circles. **c** Developed pipeline can assess aggregation severity with great accuracy and precision in both mono- and polydisperse droplet data. Shown data is applicable to experiments performed with sub-optimal concentrations of cefotaxime and kanamycin in monodisperse- and cefotaxime, chloramphenicol, ciprofloxacin, doxycycline and trimethoprim in polydisperse droplets. To assess aggregation detection performance, droplets containing aggregative bacteria were interpreted ‘positive’, whereas droplets with homogenous growth and no growth were considered ‘negative’. Figure created with biorender.com.

Computationally, texture characterisation starts with constructing a grey-level co-occurrence matrix (GLCM) to assess how often two pixels in a predefined distance and direction with specific values appear on an image. GLCM alone is not a descriptor of texture and estimating texture must be presented in a single representative value to determine one measurement for an image or an object. For this, Haralick [21] famously proposed 14 definitions. Out of these, sum variance of texture proved to be the most suitable for aggregation analysis in droplets as shown in Supplementary file 3.

Currently, the textural values of each droplet in our pipelines are classified using a strict threshold as demonstrated in Supplementary File 3. Only providing the most basic functionality of a strict threshold to classify data into bins is seen as a major limitation of Cellprofiler^™^. For more advanced classification demands, users can use a sister-software CellProfiler Analyst incorporating machine learning methods, or alternatively, employ other specialised software or programming languages with the necessary advanced capabilities. The principles and protocol for implementing the pipeline is outlined in Supplementary file 1.

In some droplets, bacteria failed to form traditionally seen large clusters and instead exhibited fine, pinpoint-like aggregation patterns. Because these micro-aggregates were very small and difficult to distinguish from homogenous droplets based on texture alone, they were classified together with the homogenous group. This limitation highlights a potential improvement area where adaptive classification approaches, such as *k*-means clustering, could help determine whether a droplet’s texture more closely resembles aggregation or homogenous growth.

Furthermore, CellProfiler^™^ is equipped to detect and exclude extraneous objects during the analysis of the objects under interest. The main analysis of target objects remains unaffected while simultaneously providing valuable information on the presence of additional objects to the user. This translates to the capability of assessing the presence of additional objects (e.g. microplastic) inside the droplets that can affect the aggregation patterns of bacteria. An easy setup of the pipeline for microplastic beads alongside bacteria in droplets is assisted with a Supplementary file 2.

In general, empirical experimentation in our laboratory has shown universality and effectiveness of our pipeline. In our experience, texture characterisation a) does not rely on defining thresholds to determine the boundaries of objects under interest (aggregates) inside droplets and b) textural properties are independent of the experiment’s nature or conditions.

The method proves to be computationally efficient due to eliminating the need for additional modules within the pipeline to perform multiple measurements. Solely analysing texture and classifying droplets to their respective categories using 10 16-bit monodisperse droplet image sets (30 images total) with a resolution of 2048×2048 pixels with 8 workers running takes 55 seconds.

Additional object detection (such as microplastics) doubles the duration of the same analysis, resulting in a processing time of exactly 2 minutes. This results mainly from a module calculating the illumination function to correct uneven lighting on the images, which can be prominent in brightfield images and can alter the detection of microplastics on an image. In case illumination correction is not needed, textural characterisation of abovementioned droplet dataset with additional object detection inside droplets is reduced to 1 minute and 16 seconds of processing time.

Calculating the texture of droplets via sum variance correlates well with aggregation in droplets as demonstrated in Figure 1b. Figure 1c demonstrates the accuracy of our pipeline to assess the severity of aggregation up to 94.2% (ranging from 93.5% to 94.8%) and precision of 80.7% (ranging from 78.2% to 82.8%) in monodisperse droplets. Similar performance could be achieved in polydisperse droplets with an accuracy of 96.9% (range = 94.0 − 98.6%) and precision of 89.3% (range = 65 − 100%).

Users must be aware that the polydisperse data may be more unpredictable - additional assessments may be valuable to determine an appropriate texture scale for analysing larger droplets in the polydisperse dataset. In datasets with high droplet size variability, larger droplets often contain larger bacterial aggregates. When a small texture scale – set according to the smallest droplets - is applied to these larger droplets, the resulting sum variance values may be underestimated. This occurs because many neighbouring pixels within large aggregates have similar intensities. Therefore, when analysing highly polydisperse datasets, users should adjust the texture scale to reflect the typical droplet size or minimize droplet size variability by filtering out droplets that are significantly smaller or larger than average.

As proof-of-concept, we quantified bacterial aggregation in different experimental scenarios as demonstrated in Figure 2a. The software was tested with droplets infused with *Escherichia coli*, antibiotics in varying concentrations and additionally, metals or microplastic beads. Further data analysis of these testing results revealed significant variations in aggregation patterns.

**Fig. 2.**
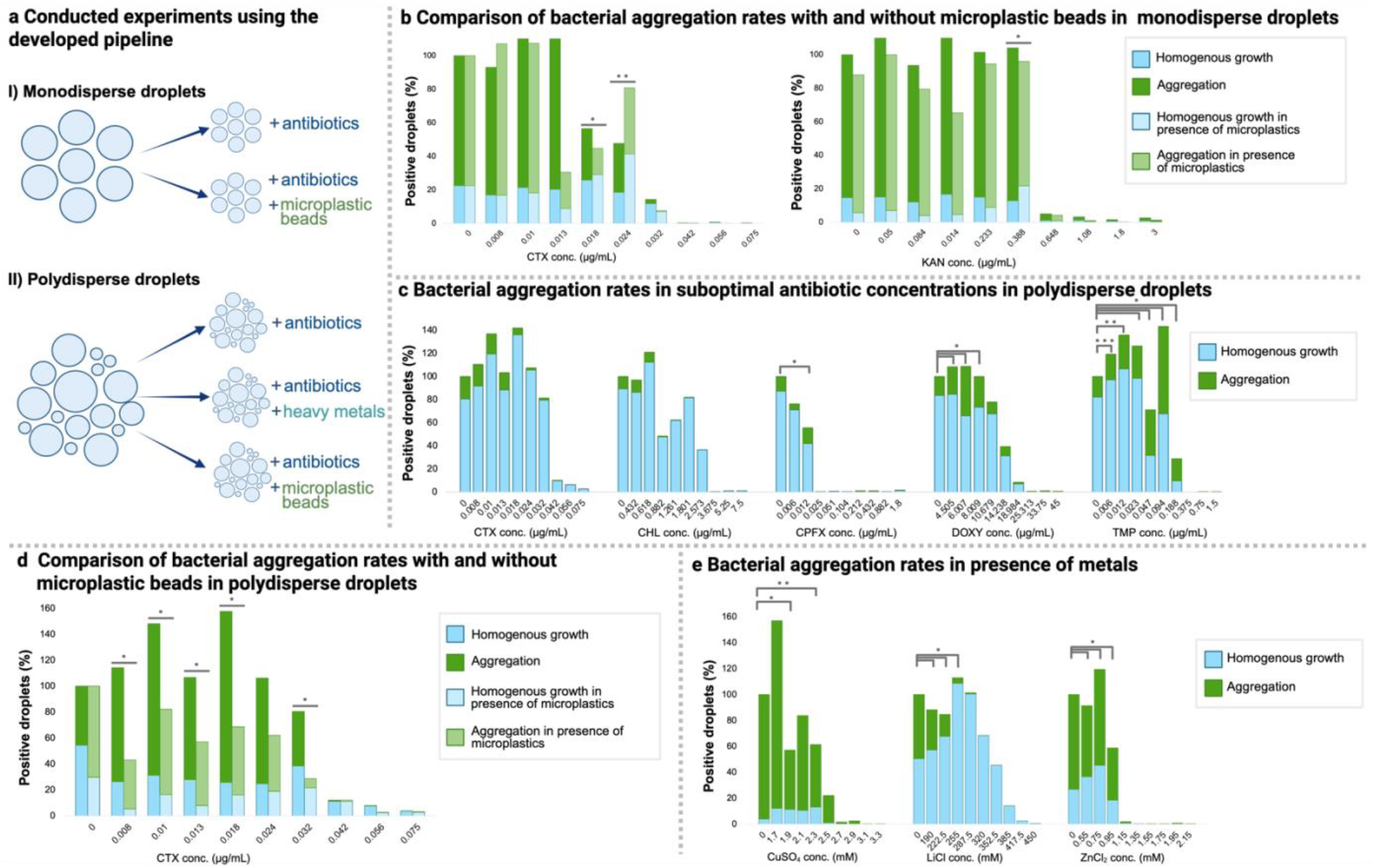
Texture-based aggregation assessment is applicable in various experimental setups. **a** The aggregation pipeline described in the current paper was applied to various datasets using *E. coli*. **b** Monodisperse droplets harbouring bacteria were infused with suboptimal concentrations of antibiotics and in an additional experiment, with microplastic beads. Microplastic beads are seen to significantly decrease aggregation where cefotaxime (CTX) concentration is 0.018 μg/mL (p<0.001), but increase aggregation at 0.024 μg/mL (p=0.026). In experiments conducted with kanamycin (KAN) with a concentration of 0.388 μg/mL, aggregation is decreased in the presence of microplastics (p<0.001). **c** In polydisperse droplets, a significant increase in aggregation compared to droplets without any antibiotics is seen in some concentrations of ciprofloxacin (CPFX at a concentration of 0.012 μg/mL, p<0.001), doxycycline (DOXY at a concentration of 4.505 μg/mL, 6.007 μg/mL, 8.009 μg/mL and 10.679 μg/mL, p<0.001) and trimethoprim (TMP at a concentration of 0.006 μg/mL, p=0.016; 0.012 μg/mL, p=0.008 and 0.023 μg/mL, 0.047 μg/mL, 0.094 μg/mL, 0.188 μg/mL, p<0.001). **d** In polydisperse droplets using parallel experiments of cefotaxime-infused droplets with and without microplastic beads, CTX at concentrations of 0.008 μg/mL, 0.01 μg/mL, 0.013 μg/mL, 0.018 μg/mL and 0.032 μg/mL decrease aggregation significantly (p<0.001). Similarly to experiments conducted with monodisperse droplets shown on Figure 2b, microplastic beads appear to have an antagonistic effect on the bacterial natural defence mechanisms against antibiotics, resulting in increased sensitivity to antibiotics. **e** Droplets containing *E. coli* and metal solutions demonstrated a significant decrease in aggregation. In experiments conducted with CuSO_4_, a reduced presence of aggregates was shown in concentrations of 1.9 mM and 2.3 mM with p values of 0.002 and <0.001, respectively. A similar effect was observed with experiments using LiCl and ZnCl_2_, where aggregation and overall viability is effectively reduced in response to increasing levels of LiCl and ZnCl_2_ (• - p<0.001). Figure created with biorender.com.

First, antibiotic exposure modulates the aggregation of bacteria depending on the antibiotic type (Figure 2c). The most pronounced levels of increased aggregation are observed in the nucleic acid synthesis targeting antibiotics CPFX and TMP. The cause of the phenomenon is likely the release of small amounts of extracellular DNA into the bacterial environment, which is well known for its aggregative characteristics [22]

A contrasting observation regarding protein targeting antibiotics was observed, where CHL did not exhibit aggregation inducing effect whereas DOXY did so significantly. This difference may be related to where the antibiotics bind to the ribosome – CHL is known to bind to the 50S A site, whereas DOXY affects the 30S subunit. In literature, other antibiotics targeting 30S subunit are known to boost bacterial aggregation up to 100-fold [23]. The insignificant results of CHL align well with previous experiments, where biofilm formation capacity was significantly downregulated in exposure to CHL [24].

Although antibiotics targeting the cell wall synthesis, such as CTX, are well known for their aggregative qualities [4], [22], our experiment did not witness a significant increase in aggregation. The mechanism driving this response has yet to be identified.

Strikingly, our data is indicating that the presence of microplastics in the bacterial environment could have a negative effect on the bacterial aggregation. A significant reduction in aggregation rates and increased sensitivity to CTX was shown in both mono- and polydisperse droplet experiments (Figure 2b and 2d). The effect is speculated to be linked to oxidative stress-inducing properties of microplastics, which could enhance antibiotic effect on bacterial cells [25][26][27]. Our preliminary findings highlight the need for more systematic studies on bacterial aggregation dynamics in the presence of microplastics, given that many studies also report microplastics as an aggregation platform and a promotor [26][28].

Our experiments revealed a significant influence of metals on bacterial aggregation patterns. As shown on Figure 2e, introduction of metals to bacterial environment rapidly decreases the aggregation levels as well as the viability of bacteria. Although CuSO_4_, LiCl and ZnCl_2_ salts are known for their innovative antimicrobial applications prospectsin biotechnology [29][30][31], they are also known to enhance antimicrobial resistance gene transfer [32][33]. Not all is known in terms of physiological response of bacteria to these substances and droplet technologies may prove valuable to further determine the cellular and molecular responses occuring within smaller subpopulations. There are some limitations to our study. First, insufficient droplet aeration may influence aggregation patterns as a clear relationship between imbalanced gas concentrations and increased aggregation is known. Although this effect could ideally be evaluated by comparing the droplet-based method with a more conventional approach assessing bacterial aggregation, the absence of standardised methods and protocols prevents such comparison.

To overcome this limitation, each experiment included a reference droplet dataset that contained no antimicrobials or pollutants. This served as a baseline for comparison. As the potential impact of poor aeration would be consistent across all experimental conditions, including those with varying antibiotic concentrations or pollutants, it is unlikely to bias our findings.

Furthermore, antibiotic leakage is a known challenge in water-in-oil droplet systems. The likelihood of diffusing into neighbouring droplets increases with smaller and more hydrophobic antibiotic molecules [34]. Leakage is both concentration- and distance-dependent, meaning that higher antibiotic concentrations and droplets in close proximity can increase cross-talk between droplets [34]. In polydisperse datasets, numerous small droplets occupying the gaps between larger ones may further facilitate diffusion, suggesting that some degree of leakage could have occurred under these conditions. However, the phenomenon is expected to extend across all experiments and therefore does not invalidate our data and our results highlight bacterial aggregation patterns in the presence of suboptimal antibiotic concentrations, microbeads, and metal pollutants. In future work, the use of thermosetting oils could be explored, as forming a cured oil matrix around droplets may help prevent leakage and cross-talk between them [35], [36].

## Conclusion

Texture based image analysis pipeline was established in a user-friendly and open-source software CellProfiler™.The method is considered a fit tool to unveil bacterial aggregation mechanisms in response to different environmental factors, such as suboptimal antibiotic concentrations and microplastic bead presence. The solution addresses the unmet need for efficient analysis of aggregated bacteria within microfluidic droplets in bulk, paving the way to improve or develop novel antimicrobial strategies.

Providing high-throughput methods for droplet image analysis bridge the gap in bioinformatics expertise among laboratory professionals, enabling the advancement of novel antimicrobial strategies and evaluating their efficacy. User-friendly approach of the method promotes the automation of image analysis in droplet microfluidics, making it nearly an instantaneous process while ensuring reproducible and quantitative results. While the current study design used bacterial aggregates as an object of interest inside droplets, texture characterisation could be applicable in various other experiments, such as cellular (platelets, tumor cell clusters) or protein aggregation experiments. Moreover, the calculation of droplet content texture values in general are easily applicable in other software (e.g. ImageJ/Fiji) or programming languages (e.g MATLAB®, Python). This ensures that the method is accessible and easy to employ regardless of the preferred tool used for image analysis.

It is recommended that further research be undertaken regarding experimentation with different tools, particularly script-based tools. This may aid in establishing a greater degree of understanding of droplet analysis in microbiology and other fields, perhaps leading to the development of advanced tools dedicated to droplet image analysis. Additionally, it would be beneficial to develop an image analysis pipeline to determine the size and shape of aggregates, which would provide further valuable insight on bacterial aggregation patterns and -behaviour influenced by various contaminants in their environment.

## Methods

### Microbiology

*E. coli* strain JEK 1036, is a wild-type *E. coli* W3310 strain labelled with green fluorescent protein [37]. To prepare *E. coli* for the study, one colony of *E. coli* grown on Luria-Bertani (L-B) agar (BioMaxima, Poland) was dissolved in 5 mL of L-B broth (BioMaxima, Poland) and incubated overnight at 37° C. After incubation, the optical density of the culture was measured. The culture was diluted with the mixture of fluorophore Alexa Fluor™ 647 (Invitrogen, Thermo Fisher Scientific Inc., USA) dye, antibiotics or metal solutions, and L-B broth (BioMaxima, Poland) to achieve 5.9×10^5^ - 2×10^6^ CFU/mL. As a result, bacterial density value of yielding roughly 50:50 ratio of positive and negative droplets.

The antibiotic concentrations were chosen based on the MIC curves of a 96-well plate method shown in Supplementary File 3. The MIC curves for metal solution concentrations were determined based on previously reported levels of heavy metal contamination in environmental hotspots, to best reflect real-world conditions of metal pollution and the co-selection of antimicrobial resistance mechanisms [38], [39], [40], [41]. The selection of microbeads content was aimed to produce at around 1-3 microbeads per droplet.

### Droplet generation

To generate monodisperse droplets sized 120 nL, a PDMS microfluidic chip was prepared and connected with appropriate sizes of PTFE tubing as described by Bartkova et al. [6]. To generate polydisperse droplets sized between 20 − 200 nL, test tubes containing oil, Alexa Fluor™ 647 and bacterial suspension were vortexed using Vortex-Genie® 2 (Scientific Industries, USA) for 5 seconds. Novec HFE 7500 fluorocarbon oil with 2% concentration of perfluoropolyether (PFPE)−poly(ethylene glycol) (PEG)−PFPE triblock surfactant was used for both mono- and polydisperse droplets.

### Imaging setup

For imaging, Countess™ (Invitrogen, Thermo Fisher Scientific Inc., USA) cell counting chamber slides were prepared with droplets. Approximately 18–20 µL of droplets with oil were pipetted into one chamber of the slide to fill the chamber. Images of droplets were captured using the LSM 900 Laser Scanning Microscope (Zeiss, Germany) equipped with ZEN 3.3 (blue edition) software with following settings: Plan-Apochromat 10x/0.45 objective, diode lasers 488 and 640 nm, DIC light, pinhole size 460 µm. In experiments using monodisperse droplets and polydisperse droplet experiments with solely antibiotics, 8-bit images were acquired. Polydisperse droplet experiments with microplastics and -metals were imaged in 16-bit format. All images were acquired at a resolution of 2048×2048 pixels and exported as TIFF files.

## Software and data analysis

All acquired images were analysed using CellProfiler™ software version 4.2.6 developed in Python environment (Broad Institute, Inc., USA) [42]. All experiments were carried out on MacBook Air (M1, 2020) with a macOS Monterey system version 12.0.1 (Apple Inc., USA). Data analysis was performed using Excel (Microsoft Inc., USA). For statistical testing, χ^2^-test was employed with a significance level set at a p-value of <0.05.

## Figure assembly

Composite figures were compiled using Biorender (Biorender.com).

## Data availability

All data generated or analysed during this project are included in this published article and its supplementary information files.

## Acknowledgements

This work was supported by Estonian research council grants PRG620 and MOBJD556, and by Tallinn University of Technology grant GFLKSB22.

## Author contributions

The author’s responsibilities were as follows: MSA, KL, DK, IS. were responsible of analytical pipeline development, performance analysis and validation; MSA, DG, TO were involved in data analysis, MSA, SB and OS participated in manuscript drafting, review and editing; FLS, KK, TO, and SB performed droplet generation and imaging, OS and SB were the principal investigators and participated in the design of the study, coordination and funding acquisition. All authors contributed to the interpretation of data and in writing and approval of the final manuscript.

## Competing interests

The authors declare no competing interests.

## Notes

### Competing Interest Statement

The authors have declared no competing interest.

### Summary of Updates

Additional discussion regarding aggregation pattern causes

