## Supplementary file 1 - Aggregation pipeline guidelines for "High-throughput bacterial aggregation analysis in droplets" (from another version of this preprint: version 2, 2025-07-22)

### Guidelines for Using and Troubleshooting Bacterial Aggregation Image Analysis Pipeline

#### 1. Scope

This document describes a method to determine bacterial aggregation severity in droplet experiments using texture characterisation utilised in an image analysis software CellProfiler™. Image analysis processes involve several steps which may overwhelm even experienced researchers. This guide is designed to help users effectively navigate and troubleshoot the pipeline introduced. The current document is based on a macOS version of the software – employing other operation systems may introduce some dissimilarities to processes described in the current document.

#### 2. Software and materials

Table 1. Software and materials necessary to run the pipeline and analyse bacterial aggregation in droplets.

| RESOURCE TYPE | SOURCE |
|---|---|
| **Software** | |
| CellProfiler™ version 4.2.6 | https://cellprofiler.org/ |
| Spreadsheet software – Microsoft Excel or any other equivalent | https://www.microsoft.com/en-us/microsoft-365/excel |
| **Materials** | |
| Bacterial aggregation analysis pipeline | This paper |
| Input folder containing sample fluorescence microscopy droplet images | This paper/user specified |
| Output folder for .csv file outputs from CellProfiler | User-specified |
| CellProfiler™ manual | https://cellprofiler-manual.s3.amazonaws.com/CellProfiler-4.2.6/index.html |

#### 3. Step-by-step analysis process

##### 3.1 Preparing for analysis

*This section aims to describe the preparatory work of the images to be analysed.*

3.1.1 Download the sample input images zip folder from the source described in *Table 1.*

3.1.2 Uncompress the folder and move the images to a desired location in the computer's file management system where they are easily accessible.

3.1.3 Create a folder for output files to a desired location in the computer's file management system. After image analysis completion in CellProfiler™, analysis results will be found in this folder.

3.1.4 In case own images are being used, make sure that all images are in the same folder for error-free batch processing.

##### 3.2 Getting started in CellProfiler™

*This section aims to describe the CellProfiler™ download, setup and the programs interface.*

3.2.1 Download and open CellProfiler™ according to developer's instructions from the source stated in *Table 1* under section *2. Software and materials*.

3.2.2 A 'Welcome to CellProfiler' window together with the CellProfiler™ program window itself will appear on Your screen as shown on **Figure 1**.

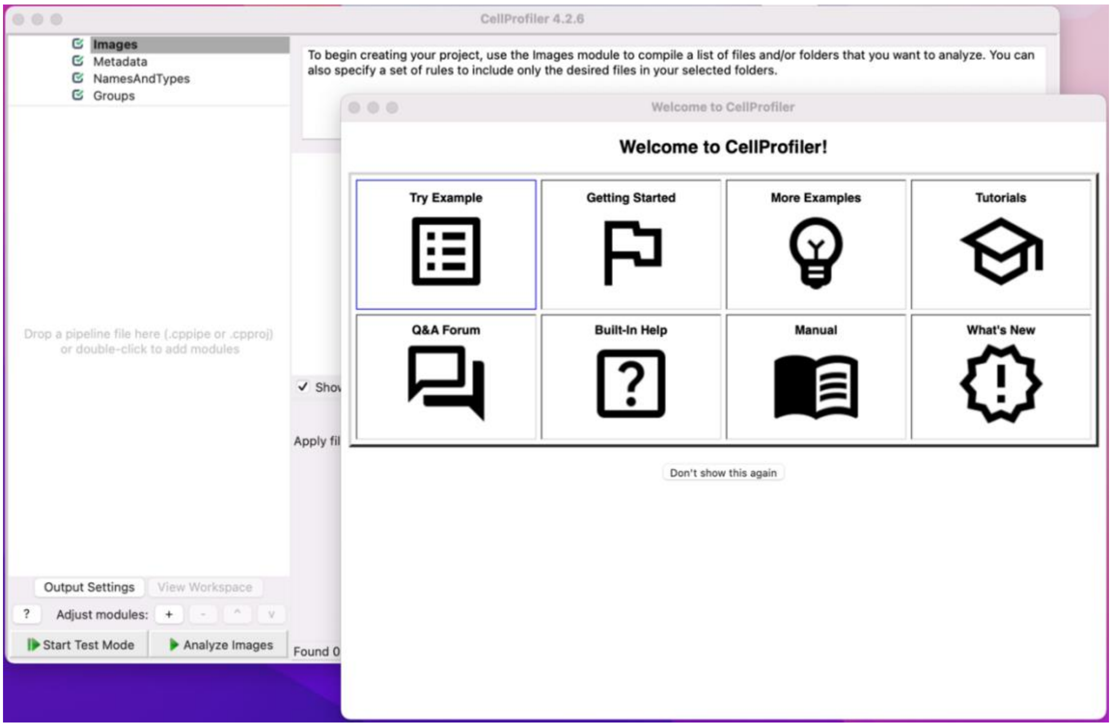


**Figure 1.** 'Welcome to CellProfiler' window and the program itself on the background.

3.2.3 The start-up window serves as a short introduction to the softwares operation principles. It is recommended to explore the information provided in the start-up page for easier use. The page serves no further functionality and the user can remove the page by clicking either ✕ or – using Windows or ⊗ or ⊖ using macOS.

3.2.4 After closing the start-up window, a full view of the program window is seen. For more convenient workflow, the window should be set in 'fullscreen' mode by clicking either ☐ using Windows OS or using macOS.

3.2.5 The program window consists of different sections with separate functionalities as described in **Figure 2**.

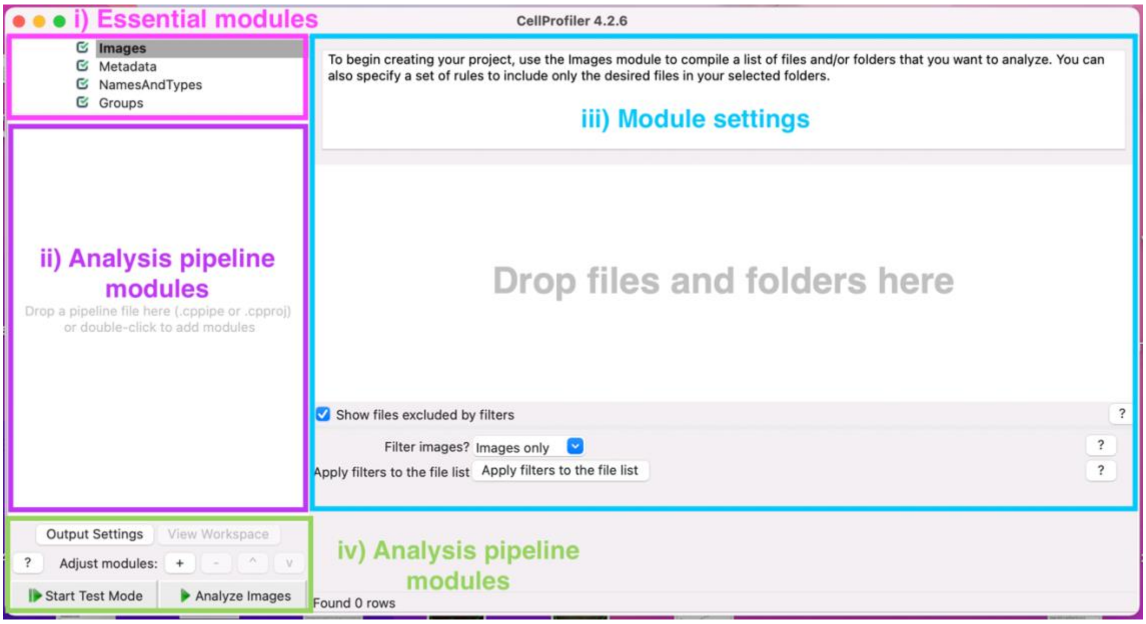


**Figure 2.** Program window functionalities. i) Section displaying essential modules for the program pipeline; ii) Section displaying modules included in the analytical pipeline specified by the user; iii) Section displaying further settings of a chosen module aka the work field; iv) Section involving different options to operate the pipeline.

👉 Whenever in doubt of the functionality of the modules setting, a question mark symbol ? located on the right side of the modules further settings section provides a short simplified explanation of the setting.

##### 3.3 Importing the pipeline

*This section aims to describe importing the pipeline* which will launch preset settings for essential modules *'Images', 'Metadata', 'NamesAndTypes'* and *'Groups'* as well as analysis pipeline modules *'IdentifyPrimaryObjects', 'MeasureObjectSizeShape', 'FilterObjects', 'MeasureTexture', 'ClassifyObjects' and 'ExportToSpreadsheet'.*

3.3.1 Find and download the pipeline from the location specified in **Table 1** under paragraph *2. Software and materials*.

3.3.2 To import the pipeline to the program window, click on ’File’ > ’Import’ > ’Pipeline from File...’ as shown on **Figure 3**.

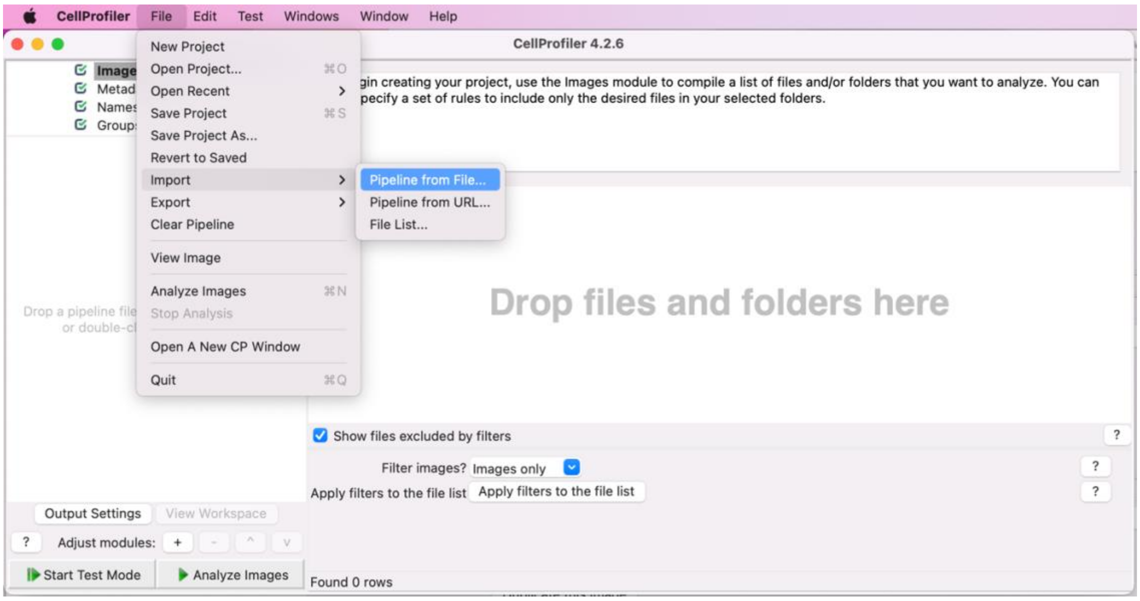


**Figure 3**. Importing a pipeline.

3.3.3 Find the pipeline from the computer's file management system file folders. Click ’Open’ in the computer’s file management system to finalise the pipeline import.

3.3.4 The following modules should appear in the program as shown on **Figure 4**.

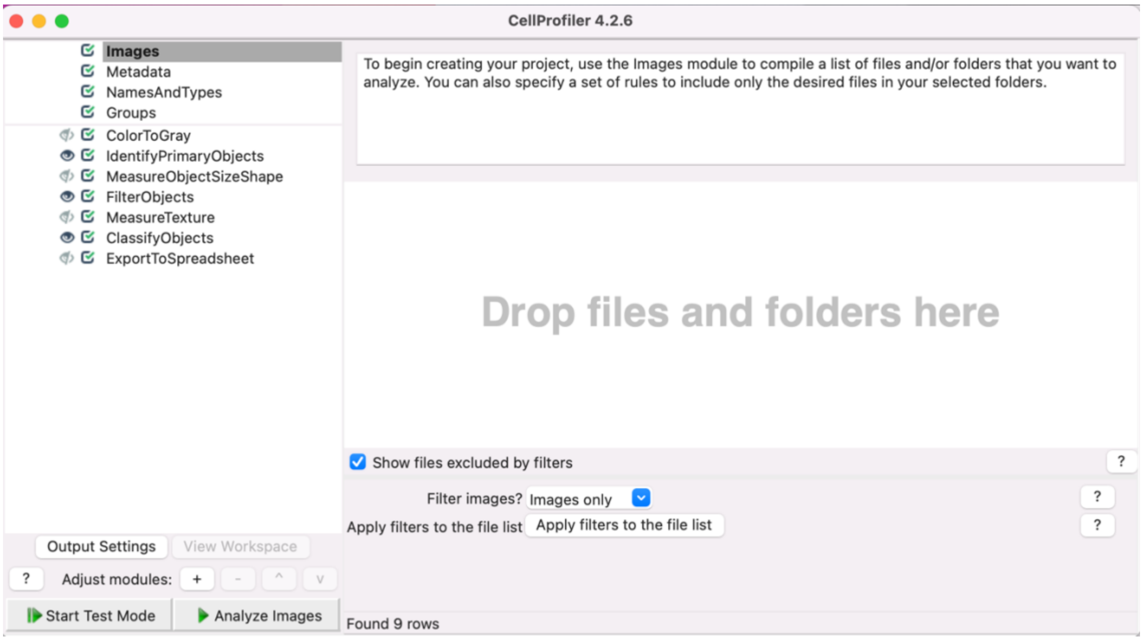


**Figure 4.** Imported pipeline. Notice the eye icon next to the analysis modules - an active eye icon will open a separate window displaying the analysis results of that module whereas an inactive eye icon does not. For smoother analysis during batch processing it is recommended to set all eye icons inactive.

##### 3.4 Essential modules setup in CellProfiler™

*This section aims to describe the setup of the essential modules* modules *'Images', 'Metadata', 'NamesAndTypes'* and *'Groups'. Find a short overview of the modules' functionalities in Table 2.*

**Table 2.** Short overview of the essential modules functionalities

| MODULE | FUNCTION |
|---|---|
| Images | Load images to analyse. |
| Metadata | Extract and associate metadata to images. |
| NamesAndTypes | Assign names to images generated during processing by which following modules will refer to images. |
| Groups | Group list of images to subsets. |

3.4.1 The first module to setup is the '**Images**' module. Click on the module.

3.4.2 Find the images folder previously prepared as described in *3.1 Preparing images for analysis.*

3.4.3 Drag and drop the images in the folder or the entire folder in the '*Drop files and folders here'* section in the program window. The setup of the 'Images' module should appear according to **Figure 5**.

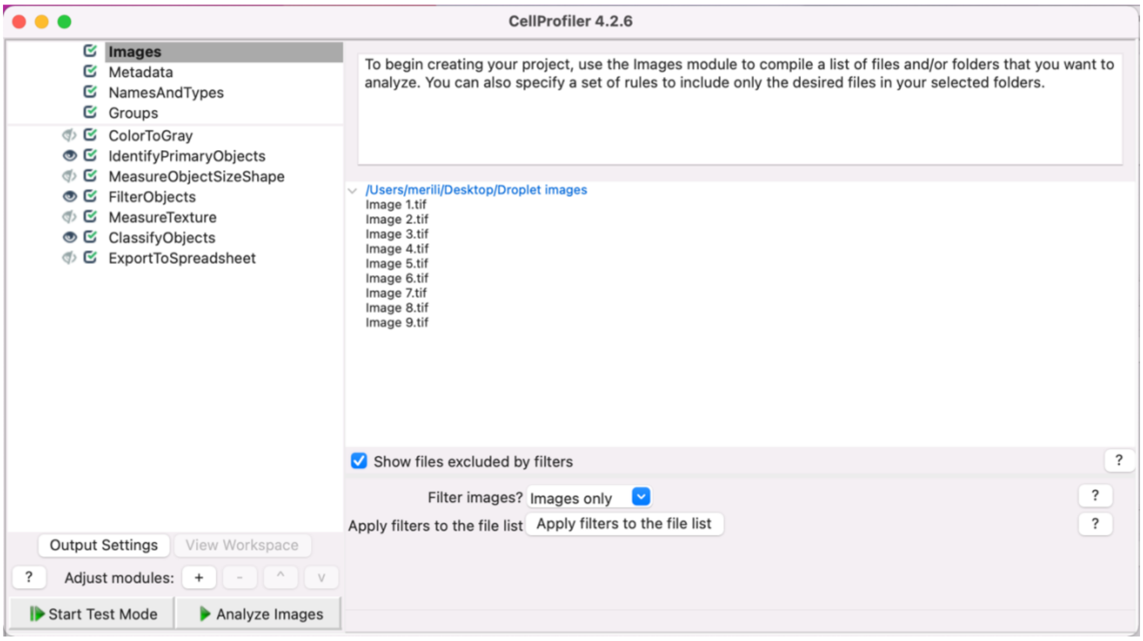


**Figure 5**. 'Images' module setup. Notice how the imported image files have appeared to the module settings section.

👉 Images module further allows to filter images in case the file list/folder imported as described in *3.4.2* contains different files from which all files will not be analysed. Different file features such as file format, file name etc could be used to filter images.

3.4.5 The second module to setup is the '**Metadata**' module. Click on the module. No metadata will be extracted. The initial settings of the 'Metadata' module should appear as shown on **Figure 6.** In case extracting metadata should become necessary, consult CellProfiler™ manual listed in section 2 of this document.

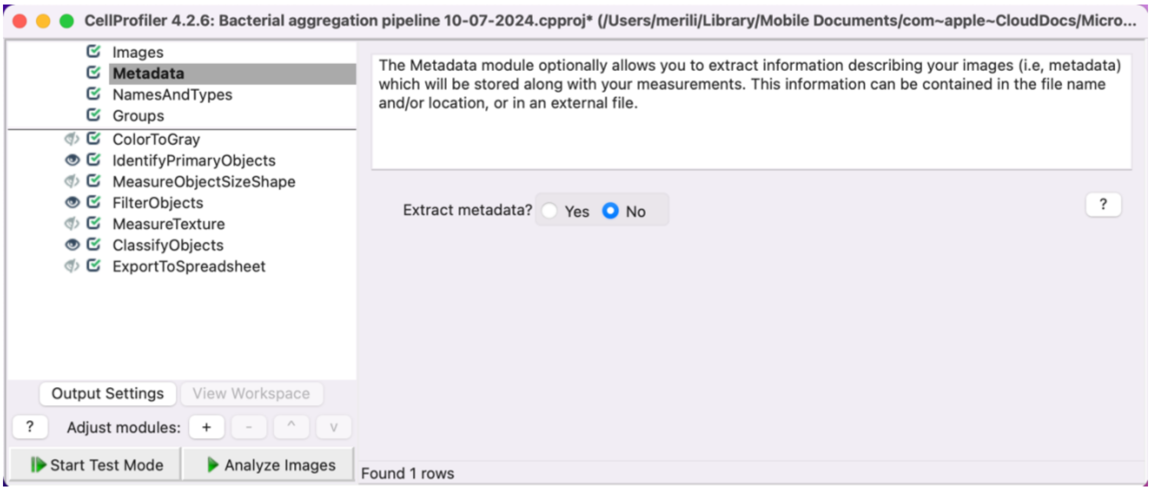


**Figure 6.** 'Metadata' module setup.

3.4.6 The third module to setup is '**NamesAndTypes**' module. Click on the module. The settings of the module 'NamesAndTypes' should appear as shown on **Figure 7.**

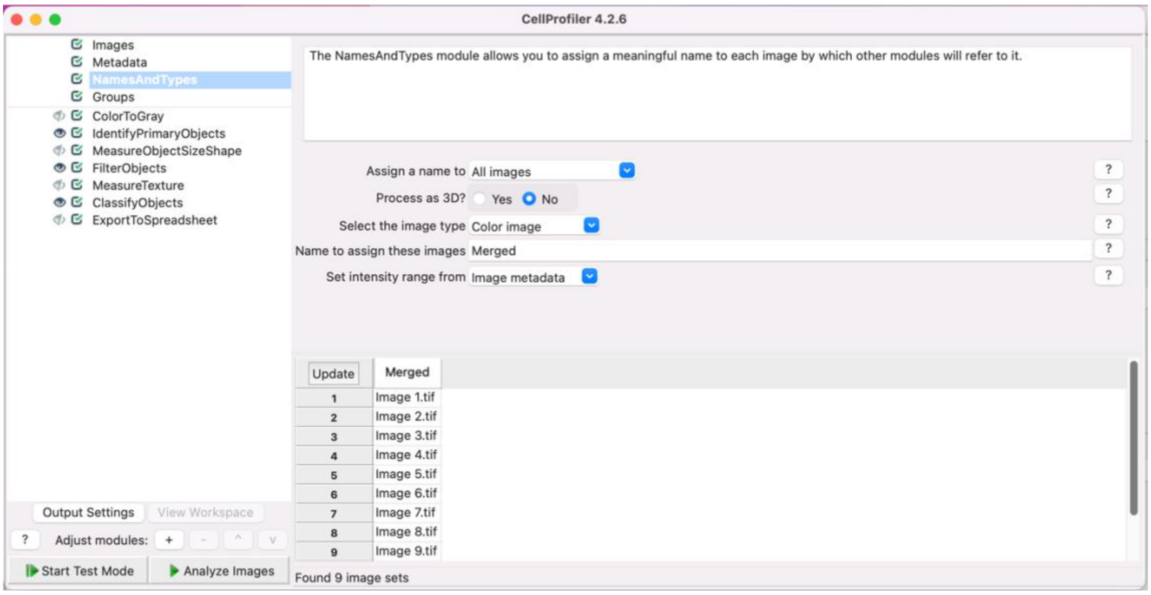


**Figure 7.** 'NamesAndTypes' module setup

3.4.7 The fourth module to setup is '**Groups**'. Click on the module. Currently, the images were not grouped as shown on **Figure 8.**

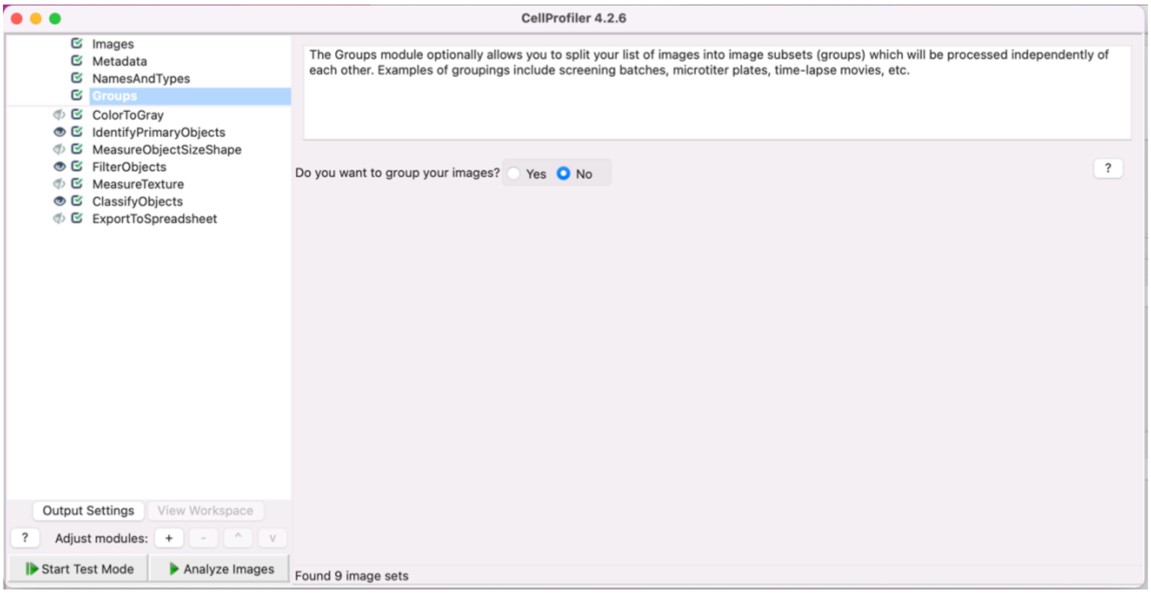


**Figure 8**. 'Groups' module setup

##### 3.5 Analysis module setup

*This section aims to describe the functionalities and setup of the modules 'ColorToGray', 'IdentifyPrimaryObjects', 'MeasureObjectSizeShape', 'FilterObjects', 'MeasureTexture', 'ClassifyObjects' and 'ExportToSpreadsheet' included in the pipeline. Find short overview of the modules' functionalities in **Table 3**.*

**Table 3**. Short overview of the modules functionalities.

| MODULE | FUNCTION |
|---|---|
| ColorToGray | Converts image with multiple colour channels to grayscale |
| IdentifyPrimaryObjects | Detects droplets using an user specified size range and adaptive thresholding strategy with Otsu method. Additionally, droplets touching the edge of the image are discarded. |
| MeasureObjectSizeShape | Measure the area and shape of identified objects. |
| FilterObjects | Discards objects based on the measurements of MeasureObjectSizeShape. Droplets of odd eccentricity or solidity are removed. |
| MeasureTexture | Measures texture of objects (droplets) detected in previous modules. Each pixel inside objects is compared to the pixel located 10 pixels horizontally to the right of the initial pixel on the image. |

| | |
|---|---|
| ClassifyObjects | Classifies objects into bins based on MeasureTexture measurements and user-set limits for the bins. Images are classified into bins using sum variance values. |
| ExportToSpreadsheet | Exports any metadata and measurements specified by the user to a .csv file. |

3.5.1 The first module to setup is ’**ColorToGray**’ module. Click on the module. The settings of the module 'ColorToGray' should appear as shown on **Figure 9**.

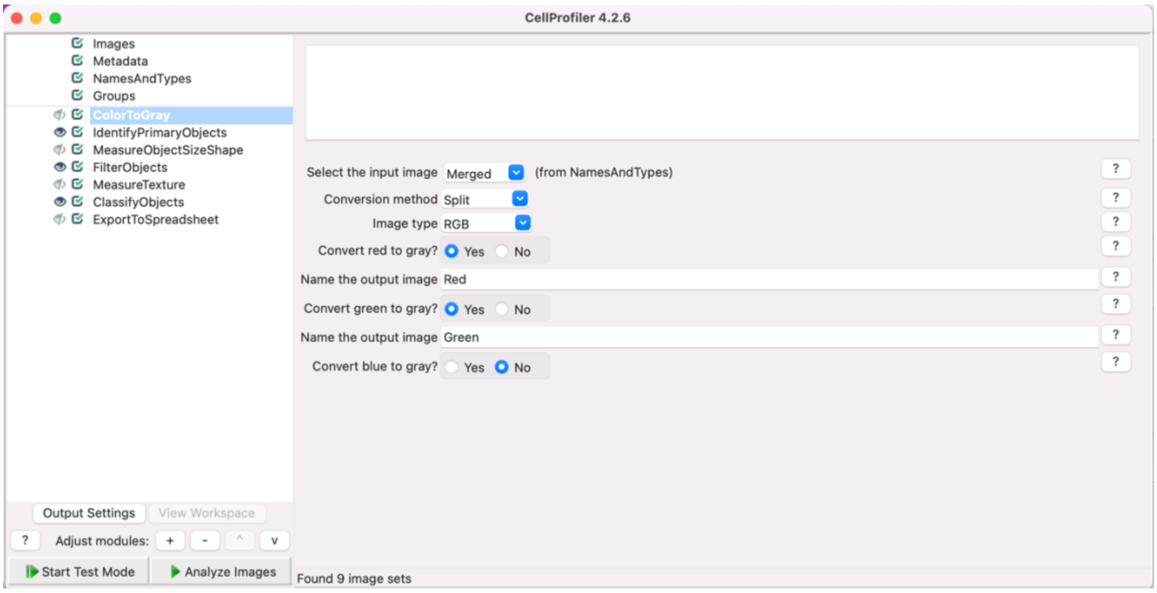


**Figure 9.** 'ColorToGray' module setup

3.5.2 The second module to setup is ’**IdentifyPrimaryObjects**’ module. Click on the module.

3.5.3 The key setting for successful droplet detection is 'Typical diameter of objects to be identified'. In case other images than provided with this paper are used, evaluate the sizes of droplets to be analysed in pixel units.

3.5.4 To evaluate droplet size in pixel units, click on the 'Images' module and on an image file. This will open the image in a separate window which includes a measurement tool.

3.5.6 Click on the measurement tool and drag a line across the droplet to measure. The measurement will be displayed in the lower right corner of the image window as shown in **Figure 10.**

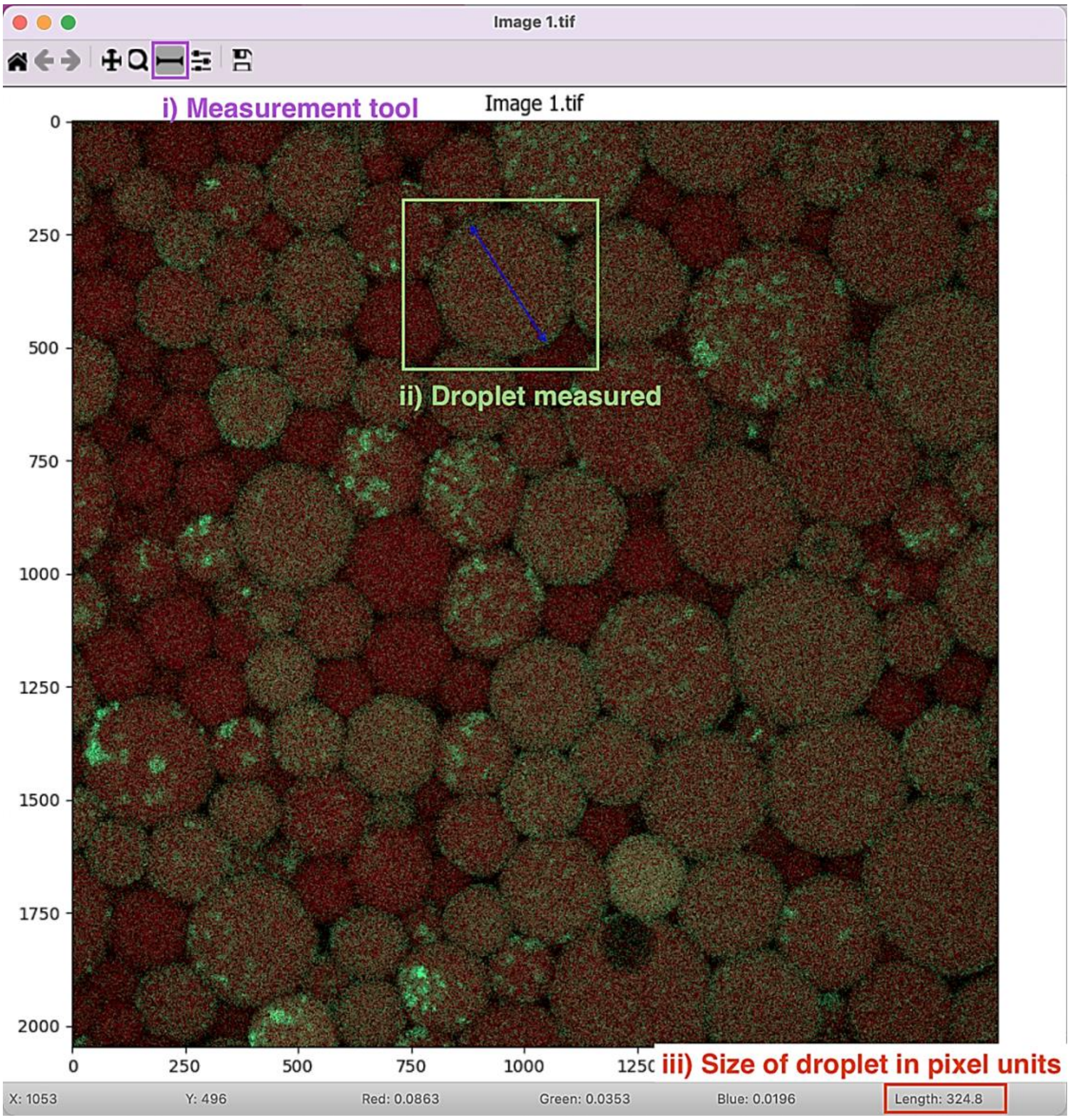


**Figure 10**. Measurement of a droplet.

3.5.5 Measure the smallest and the largest droplet on the image. These values will act as an input for the 'Typical diameter of objects in pixel units' in the 'IdentifyPrimaryObjects' module.

3.5.6 The initial settings of the 'IdentifyPrimaryObjects' module should appear as shown on **Figure 11.**

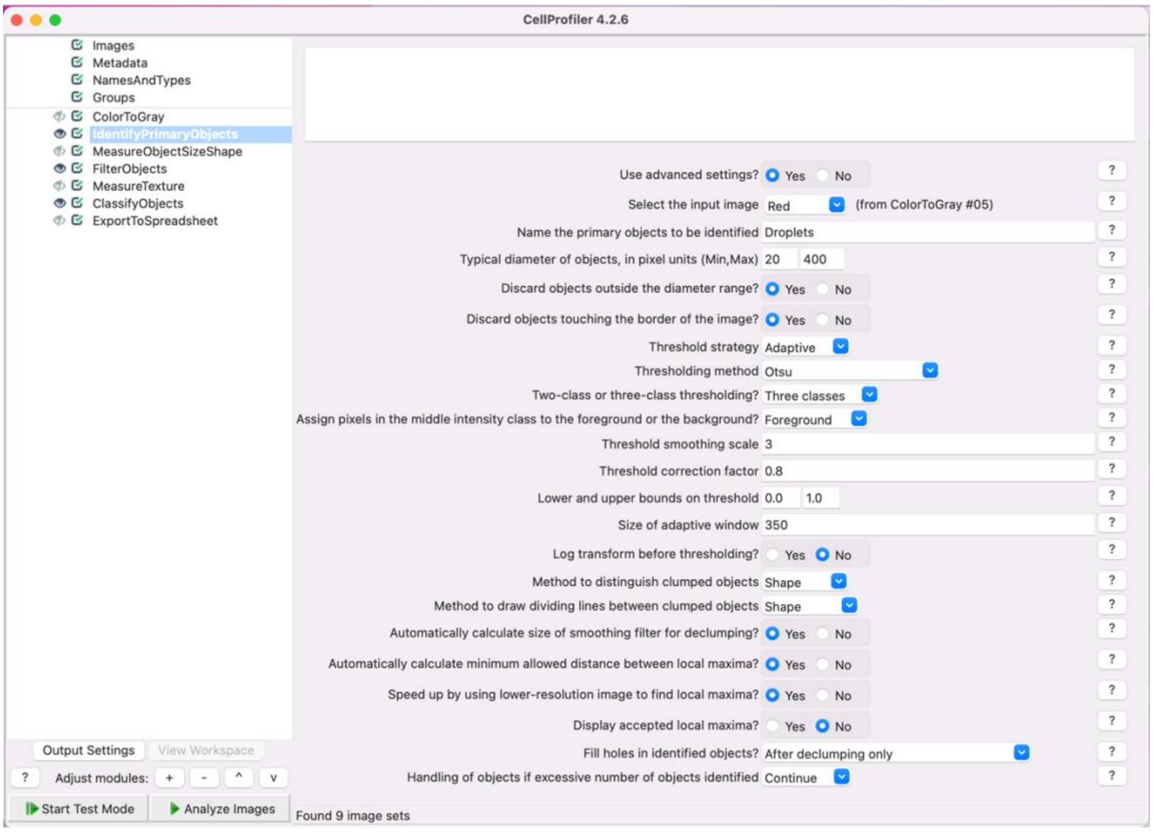


**Figure 11.** 'IdentifyPrimaryObjects' module setup. In case typical diameter of objects to be identified is changed, one should also check whether the size of the adaptive window remains appropriate.

3.5.7 The third module to setup is ’**MeasureObjectSizeShape**’ module. Click on the module. The settings of the module should appear as shown on **Figure 12.**

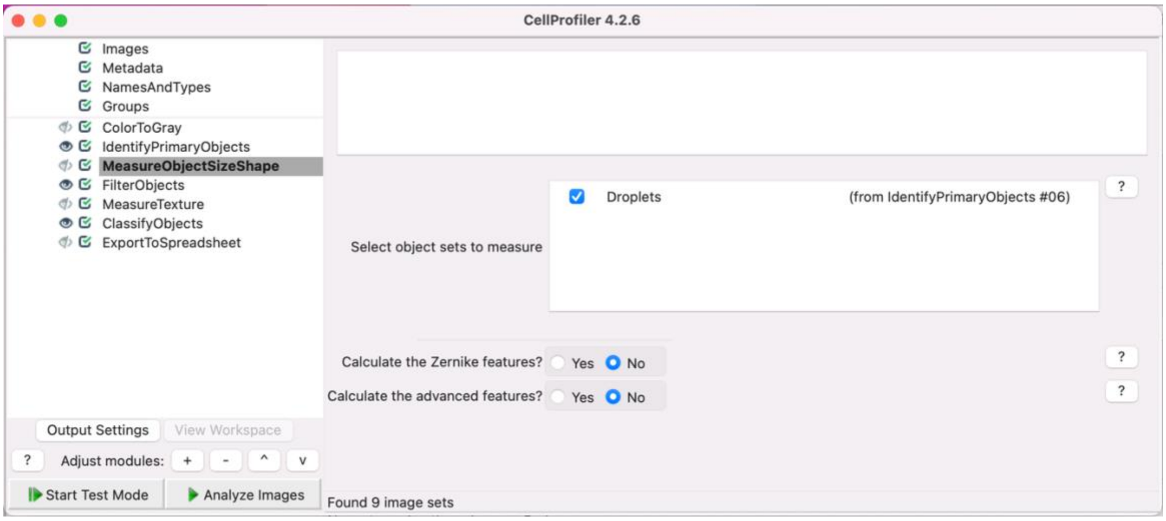


**Figure 12.** 'MeasureObjectsSizeShape' module setup.

3.5.8 The fourth module to setup is '**FilterObjects'** module. Click on the module. The settings of the module should appear as shown on **Figure 13.**

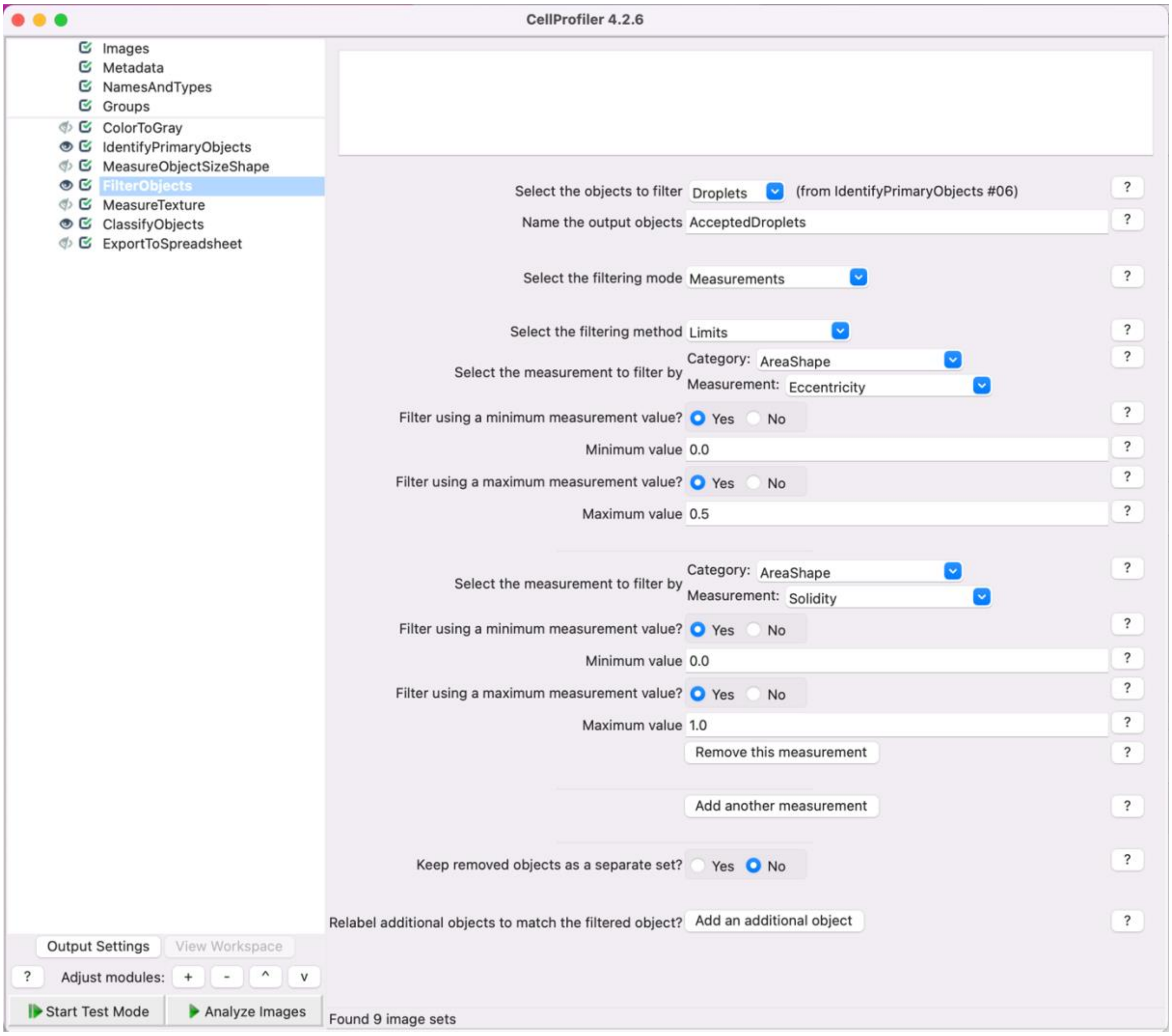


**Figure 13.** 'FilterObjects' module setup.

3.5.9 The fifth module to setup is '**MeasureTexture**' module. Click on the module.

3.5.10 Setting 'Select images to measure' is set as 'Green' as only the green channel of the image is needed for the measurement. 'Select objects to measure' should be set as 'AcceptedDroplets' which refers to droplets which were filtered out in the 'FilterObjects' module.

3.5.11 'Texture scale to measure' is currently set at 10 pixels. Texture scale should be smaller than the 'Typical diameter of objects in pixel units' specified in the 'IdentifyPrimaryObjects' module (refer to points *3.5.2* to *3.5.6*).

3.5.12 The initial settings of the 'IdentifyPrimaryObjects' module should appear as shown on **Figure 14.**

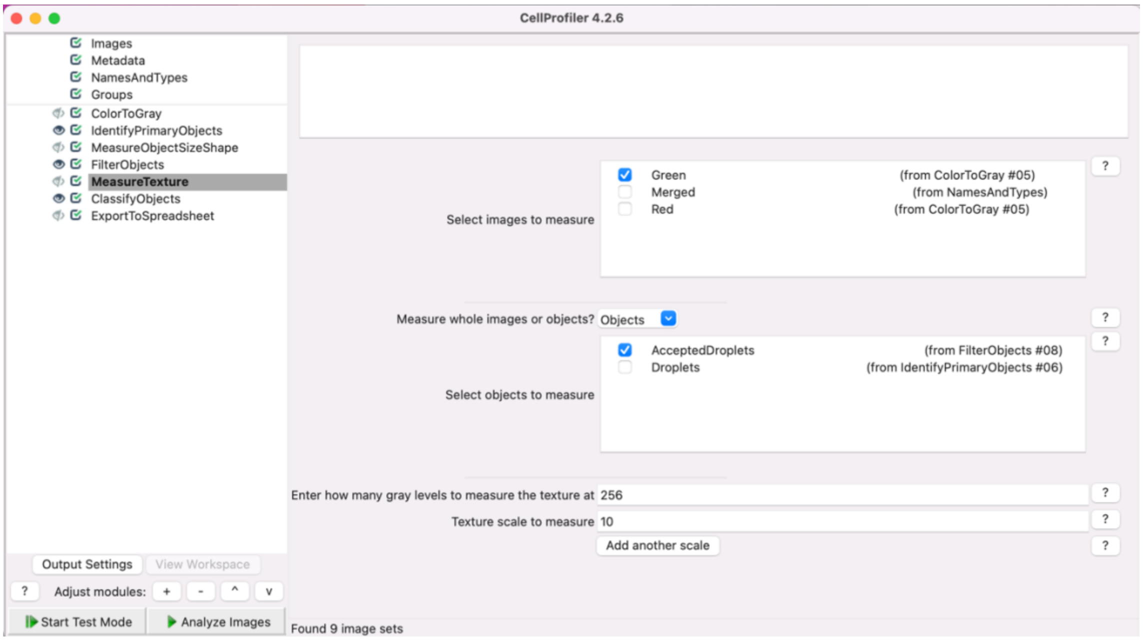


**Figure 14.** 'MeasureTexture' module setup.

3.5.13 The sixth module to setup is '**ClassifyObjects'** module. Click on the module.

3.5.14 The key setting for successful droplet classification according to their aggregation level is 'Enter the custom thresholds separating the values between bins'. Texture values may vary across different datasets depending on the image quality and other features. In case a different dataset is used other than the one provided, it is highly recommended to determine the threshold values using the Test Menu. Please consult CellProfiler™ manual listed in section 2 of this document for furhter guidance on using the Test Menu.

3.5.15 The initial settings of the 'ClassifyObjects' module should appear as shown on **Figure 15.**

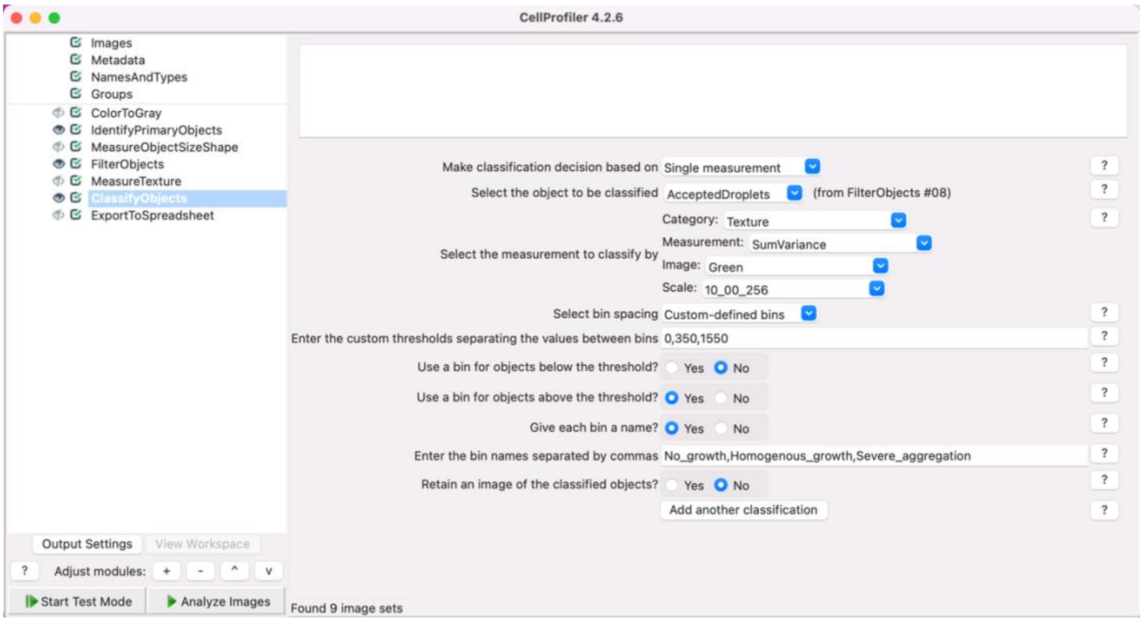


**Figure 15.** 'ClassifyObjects' module setup.

3.5.16 The last module to setup is 'ExportToSpreadsheet' module. Click on the module.

3.5.17 Set 'Output file location' as 'Elsewhere' and click on the 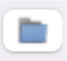 icon. Find the output folder previously prepared as described in *3.1.3,* click on the folder and follow by clicking 'Open' in the computer's file management system to finalise the action.

3.5.18 To manage the measurements exported into the .csv file, click on 'Press button to select measurements'. The key measurement to assess aggregation to export can be found under 'AcceptedDroplets' > 'Texture' > 'SumVariance'. Check the box located before 'SumVariance' as shown on **Figure 16**.

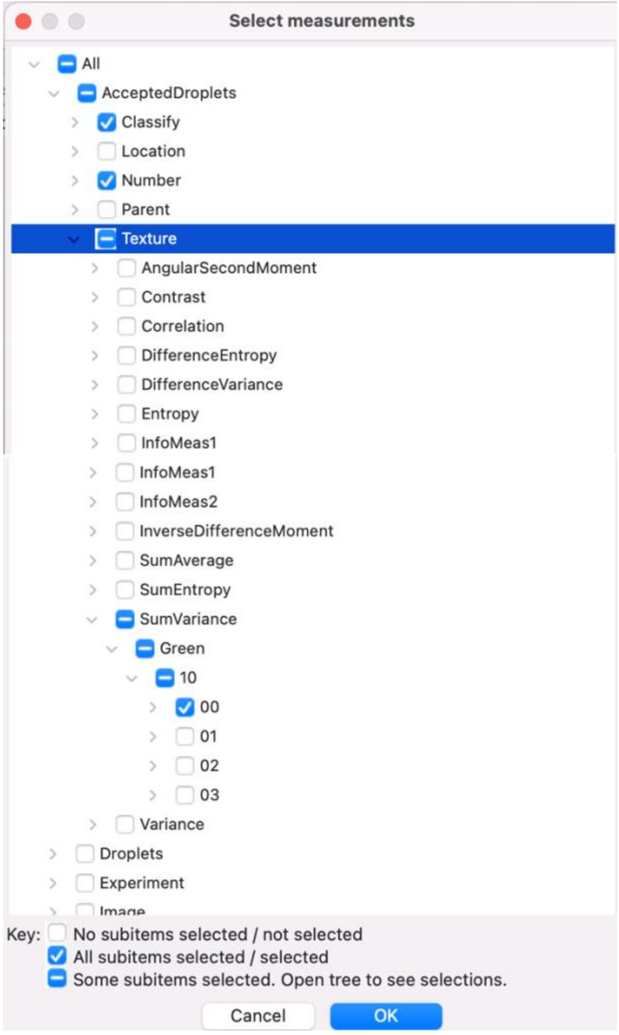


**Figure 16.** Measurement selection in 'ExportToSpreadsheet' module.

3.5.19 After selecting all needed measurements, click 'Ok'. The settings of the module should appear as shown on **Figure 17.**

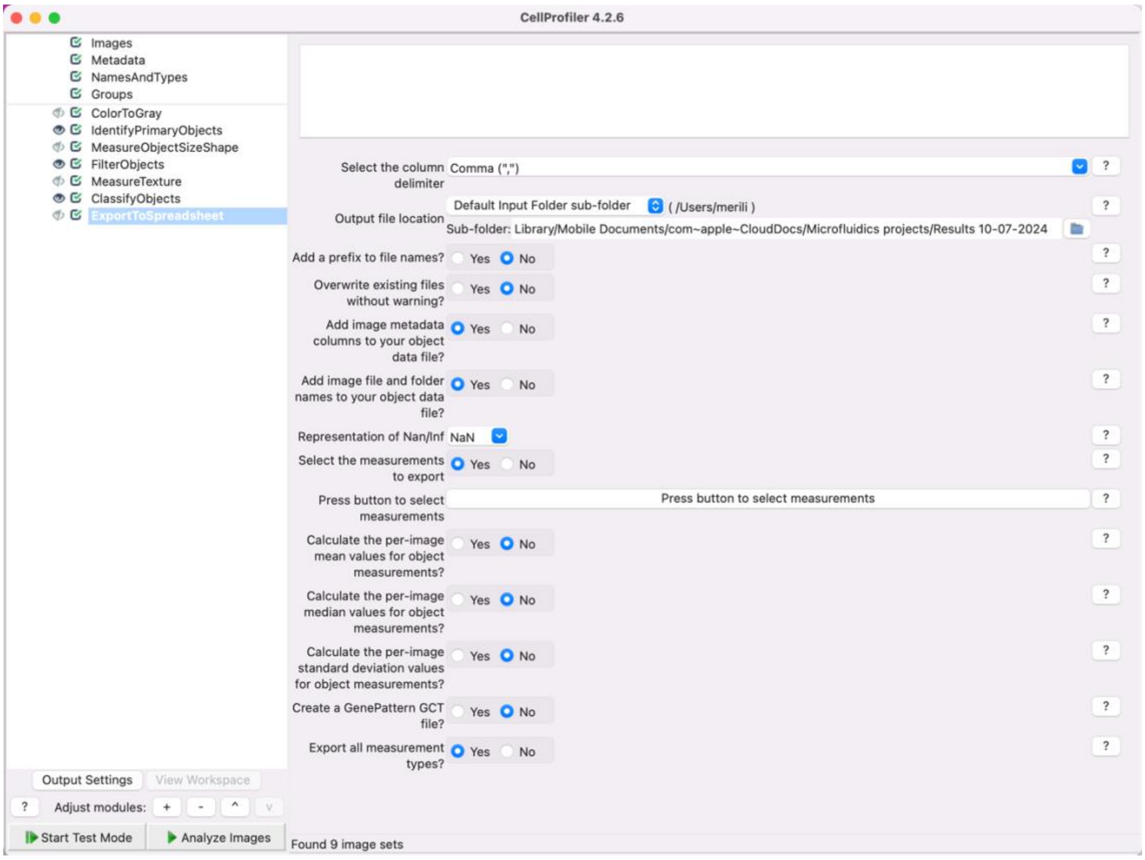


**Figure 17.** 'ExportToSpreadsheet' module setup.

3.5.19 After selecting all needed measurements, click on 'Analyze images'. For 1 image, the analysis should take around 20 seconds.

3.5.20 Find the output folder in the computer's file management system. Open the .csv file with a spreadsheet software (MS Excel) described in *Table 1* under *2. Software and materials*. Save the .csv file as an .xlsx file - performing further analysis in a .csv file format may result in data loss.

#### 4. Troubleshooting

*This section aims to describe some of the more common errors while running the pipeline presented in this guideline.*

##### 4.1 General principles of troubleshooting in CellProfiler™

4.1.1 The first indicator of an error is a red cross appeared in front of any of the modules as shown in **Figure 18**. Sliding over to the red cross (without clicking on the module) will launch a small window describing the nature of the error.

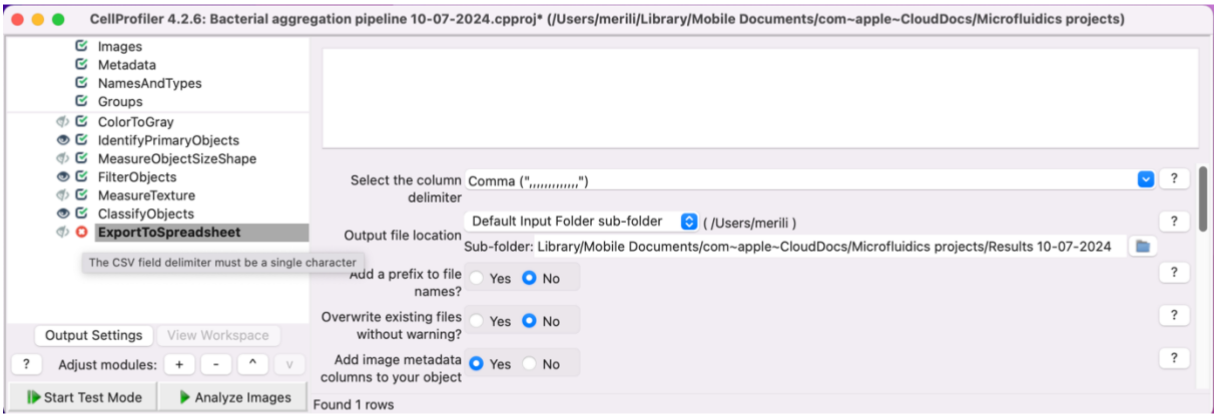


**Figure 18.** A red cross and an explanatory window describing the nature of the error in the 'ExportToSpreadsheet' module. In the current example, the setting 'Select the column delimiter' is falsely determined as 'Comma (",,,,,,,,,,,,,")'. Sliding over the red cross, an explanatory window states that 'The CSV field delimiter must be a single character'. This indicates that the setting 'Select the column delimiter' should be determined as 'Comma(",")'.

4.1.2 Whenever in doubt of the functionality of the modules setting, a question mark symbol located on the right side of the modules further settings section provides a short simplified explanation of the setting.

4.1.3 A helpful tool in troubleshooting the pipeline errors is an image.sc user community forum dedicated to CellProfiler. Find the link to the forum here.

##### 4.2 Analysis pipeline troubleshooting in CellProfiler™

4.2.1 In case the issue is of analytical nature, please consult **Table 4**.

**Table 4.** Overview of solutions for common errors regarding bacterial aggregation assessment in CellProfiler™.

| PROBLEM | SOLUTION |
|---|---|
| **IdentifyPrimaryObjects** | |
| No droplets detected | 1. Check whether the input image name in ’IdentifyPrimaryObjects’ module is set as the name of the output image assigned to red channel in ’ColorToGray’ module;<br>2. Adjust the minimum and maximum values of typical object diameter in pixel values according to object size. Object |

| | |
|---|---|
| | diameter can be measured according to paragraphs 3.5.3 and 3.5.5 of this document. |
| Background is detected as a droplet | 1. Assign pixels in the middle intensity class to the background;<br>2. Increase lower bound on threshold. |
| Undersegmentation – droplets are merged together | 1. Decrease the minimum value of typical object diameter in pixel values;<br>2. Decrease threshold smoothing scale:<br>3. Decrease threshold correction factor. |
| Oversegmentation – droplets are segmented or debris/artifacts is detected as droplets | 1. Increase the minimum value of typical object diameter in pixel values;<br>2. Increase threshold smoothing scale;<br>3. Increase threshold correction factor. |
| **MeasureObjectSizeShape** | |
| No measurements are made | Check if object sets to measure is set as the name of the primary objects to be identified in ’IdentifyPrimaryObjects’ module. |
| **FilterObjects** | |
| No filtering of the droplets | Check if selected objects to filter is set as the name of the primary objects to be identified in ’IdentifyPrimaryObjects’ module. |
| Underfiltering - unwanted droplets are not filtered out | 1. Decrease the maximum value of eccentricity;<br>2. Increase the minimum value of solidity;<br>3. In case unwanted droplets persist within droplet set suitable for analysis, consider adding another measurement to the module or a different approach for filtering. Consult CellProfiler™ manual listed in section 2 of this document. |
| Overfiltering - droplets suitable for analysis are filtered out | 1. Increase the maximum value of eccentricity;<br>2. Decrease the minimum value of solidity; |

| | |
|---|---|
| | 3. In case droplets suitable for analysis are continuously filtered out, consider removing a measurement (eccentricity/solidity) or a different approach for filtering. Consult CellProfiler™ manual listed in section 2 of this document. |
| **MeasureTexture** | |
| Texture measurements result in undefined values | Check if texture scale to measure is set a smaller value than the minimum value of typical object diameter in pixel values in ’IdentifyPrimaryObjects’ module. |
| Texture measurements of droplets show no variation | 1. Check if images to measure is set as the name of the output image assigned to green channel in ’ColorToGray’ module;<br>2. Check whether the texture scale is appropriate for the droplet size - larger droplets may need larger texture scale;<br>3. In case the error persists, it may be possible that another approach for image analysis may be appropriate. |
| **ClassifyObjects** | |
| Droplets are not classified into bins or droplets are classified into the same bin | 1. Make sure the threshold values set are appropriate for the measurement. To check the measurement values, enter Test Menu as specified in paragraph 3.5.14;<br>2. If using custom-defined bins, check whether different thresholds are separated with commas. |
| Unfiltered droplets are included in classification | Check if object to be classified is set as the name of the output objects specified in the ’FilterObjects’ module |
| **ExportToSpreadsheet** | |
| Data is not exported/the file location is unknown | Check the output file location. If necessary, create a new folder in the computers file management system dedicated to output files by choosing ’Elsewhere’ and a blue folder icon to create a new path for the output files location. |
| Data is missing from the output files | 1. Press button to select measurements and ensure there is a blue checkmark in front of the measurement of interest; |

| | |
|---|---|
| | 2. Check all output files for the missing data. |
| **Other issues not specified above** | |
| Other | 1. Consult CellProfiler™ manual listed in section 2 of this document;<br>2. Check paragraph 4.1 of this document;<br>3. Check whether the hardware of the computer used to run CellProfiler™ is appropriate. Older systems with low-performing hardware may cause errors, the software to crash/freeze or exceedingly long run times;<br>4. In case the error persists, it may be possible that another approach for image analysis may be appropriate. |
