## Supplementary file 3 - supporting data for "High-throughput bacterial aggregation analysis in droplets" (from another version of this preprint: version 2, 2025-07-22)

### Supplementary Information

**Figure S1.** Comparative suitability of textural characterisation apporach for aggregation evaluation in polydisperse droplet experiments. Five textural measurements were deemed possibly appropriate for aggregation assessment - angular second moment, inverse difference moment, variance, difference variance and sum variance. Reference set used in the polydisperse performance assessment with cefotaxime, chloramphenicol, ciprofloxacin, doxycycline, and trimethoprim median, quartiles, and min-max values of droplet's textural measurement values mentioned above were grouped according to the expert panel categorisation. Difference variance, angular second moment and inverse difference exhibit a clear contrast between textural values of droplets with no bacterial growth and bacterial growth (homogenenous and aggregated). However, the distinction capability between homogenous and aggregated droplets remain poor. A better contrast to differentiate between homogenous and aggregated droplets is seen in variance and sum variance plots, therefore, sum variance was chosen as the appropriate textural feature to distinguish aggregation from negative-and homogenous growth droplets.

**a** Cefotaxime reference set

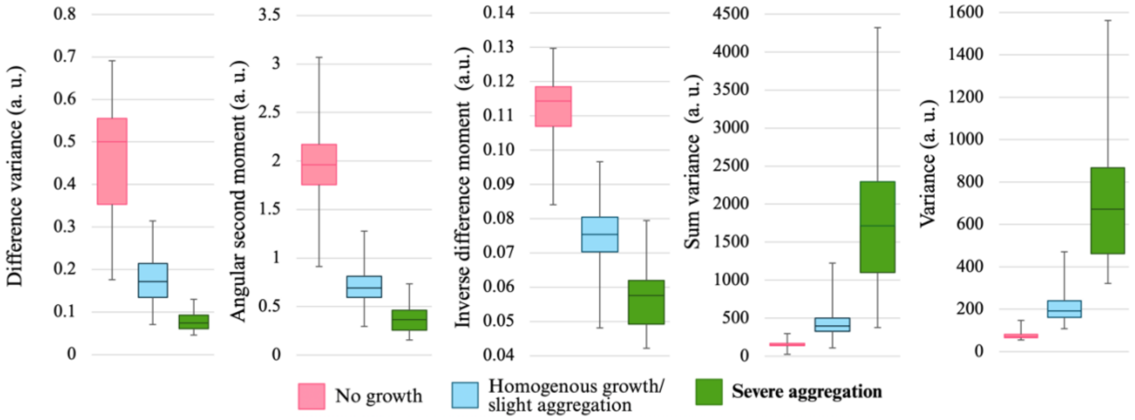


**b** Chloramphenicol reference set

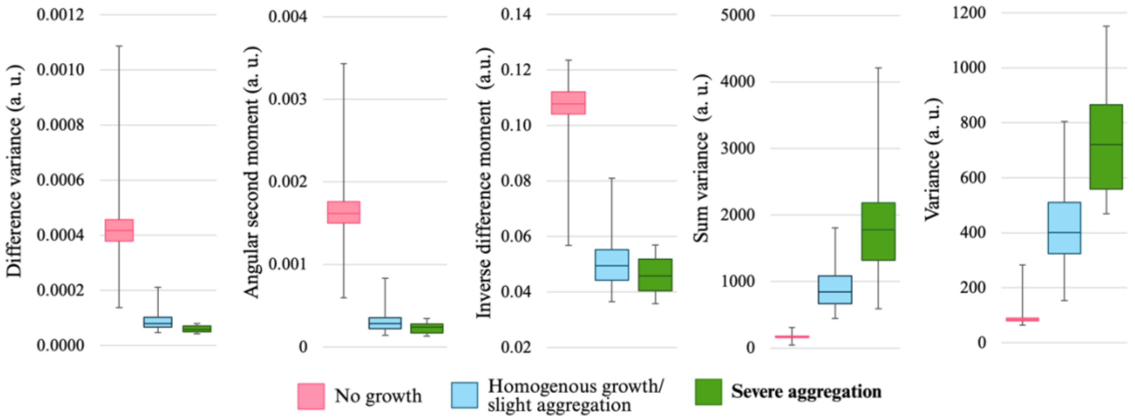

**c** Ciprofloxacin reference set

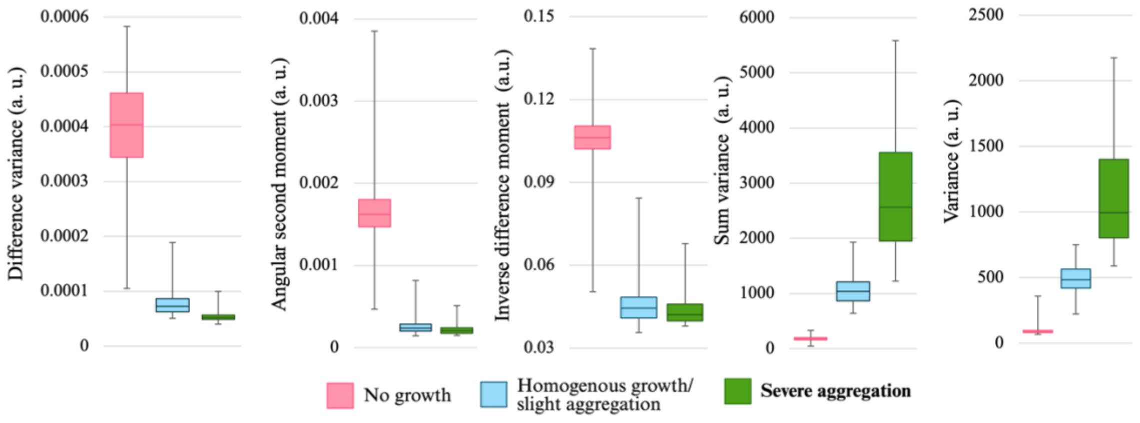


**d** Doxycycline reference set

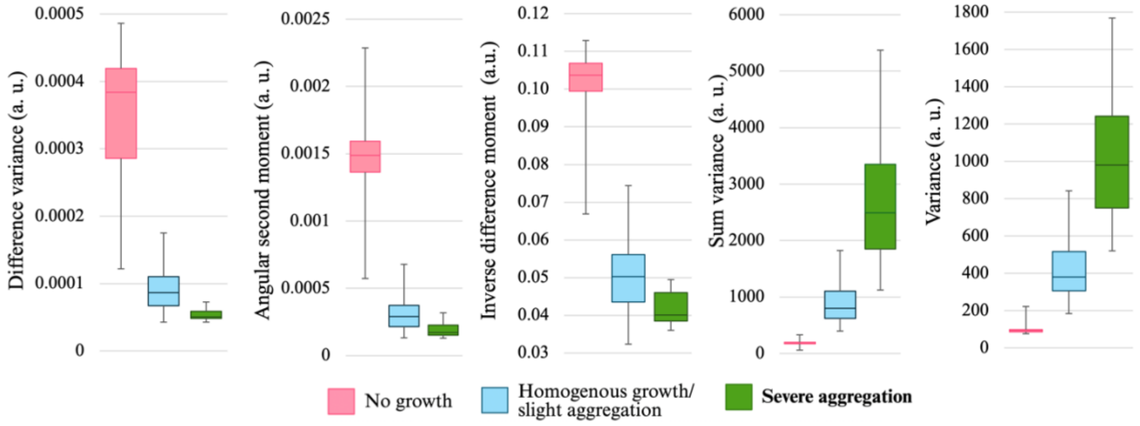


**e** Trimethorpim reference set

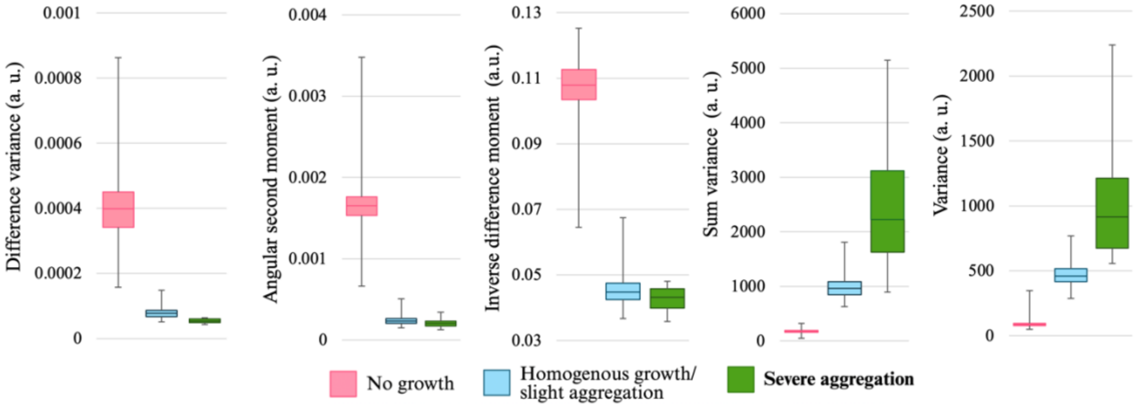

**Figure S2.** Classification thresholds for different datasets**.** To determine a classification threshold for droplet classification, sum variance values of reference sets were plotted to identify possible leaps of sum variance values within the dataset. Additionally, chosen images of each dataset with sum variance values displayed on droplets were visually assessed to further verify the appropriate classification thresholds.

**a** Monodisperse droplet experiment comparing bacterial aggregation rates with and without microplastic presence - classification thresholds for cefotaxime reference set dataset (cefotaxime conc. = 0.0 μg/mL)

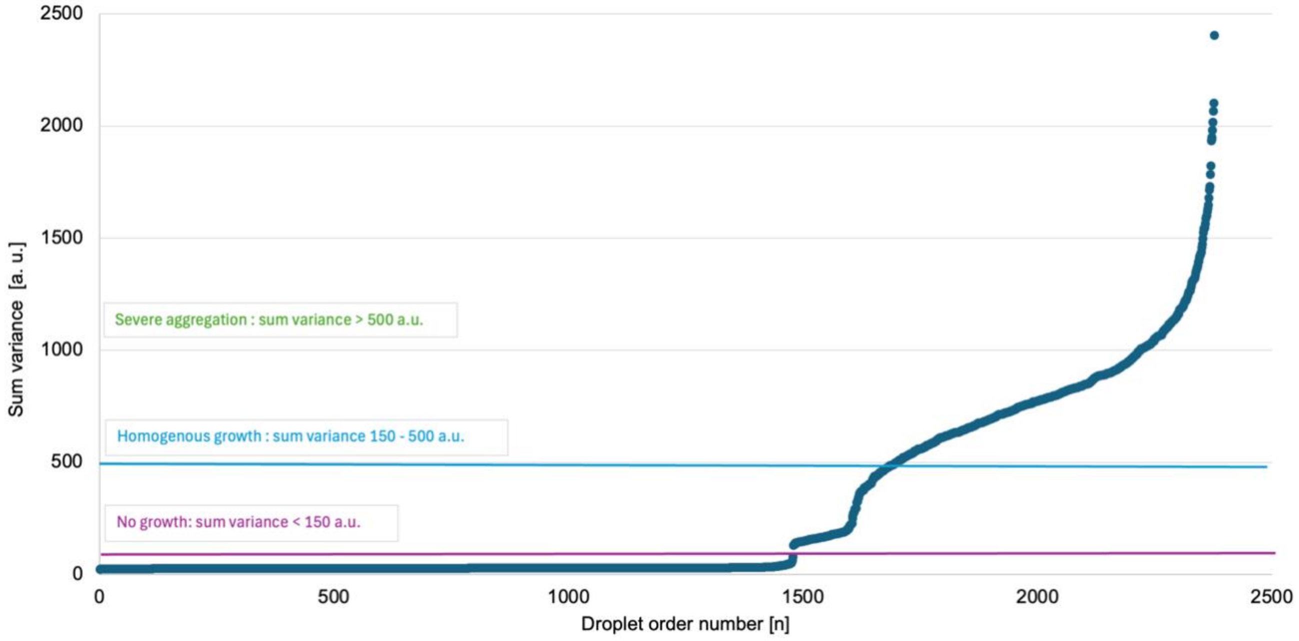


**b** Monodisperse droplet experiment comparing bacterial aggregation rates with and without microplastic presence - classification thresholds for cefotaxime dataset (cefotaxime conc. = 0.008 – 0.075 μg/mL)

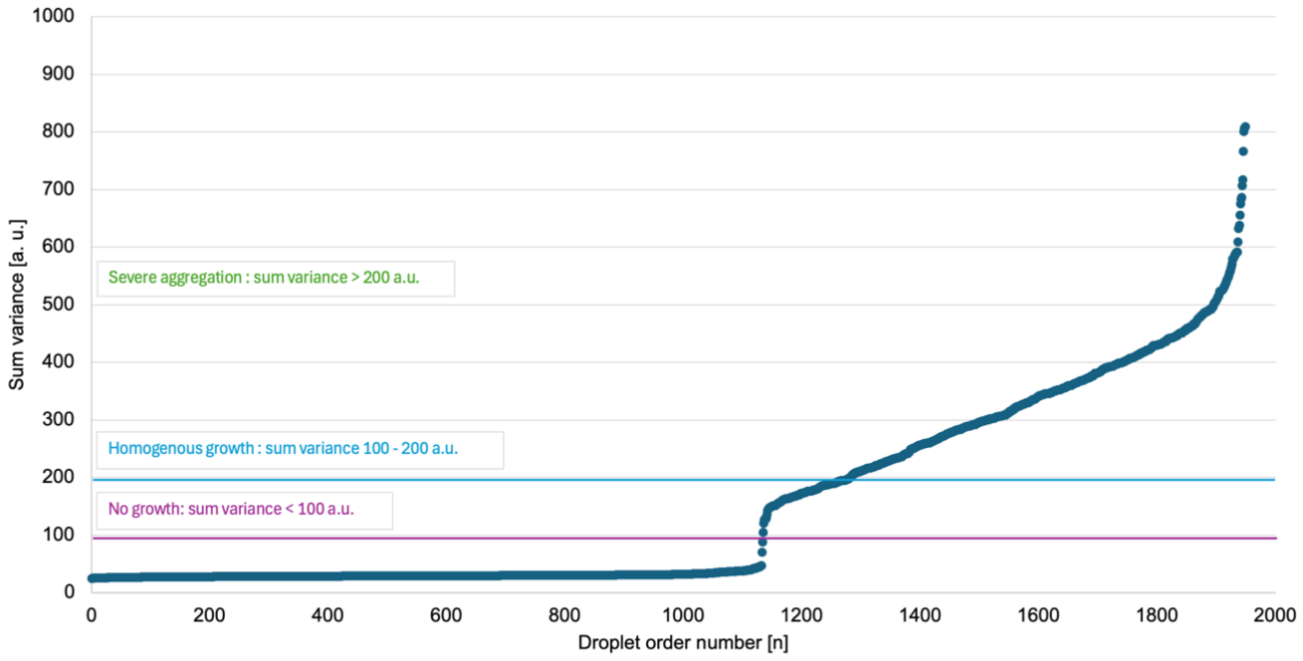

**c** Monodisperse droplet experiment comparing bacterial aggregation rates with and without microplastic presence - classification thresholds for kanamycin dataset

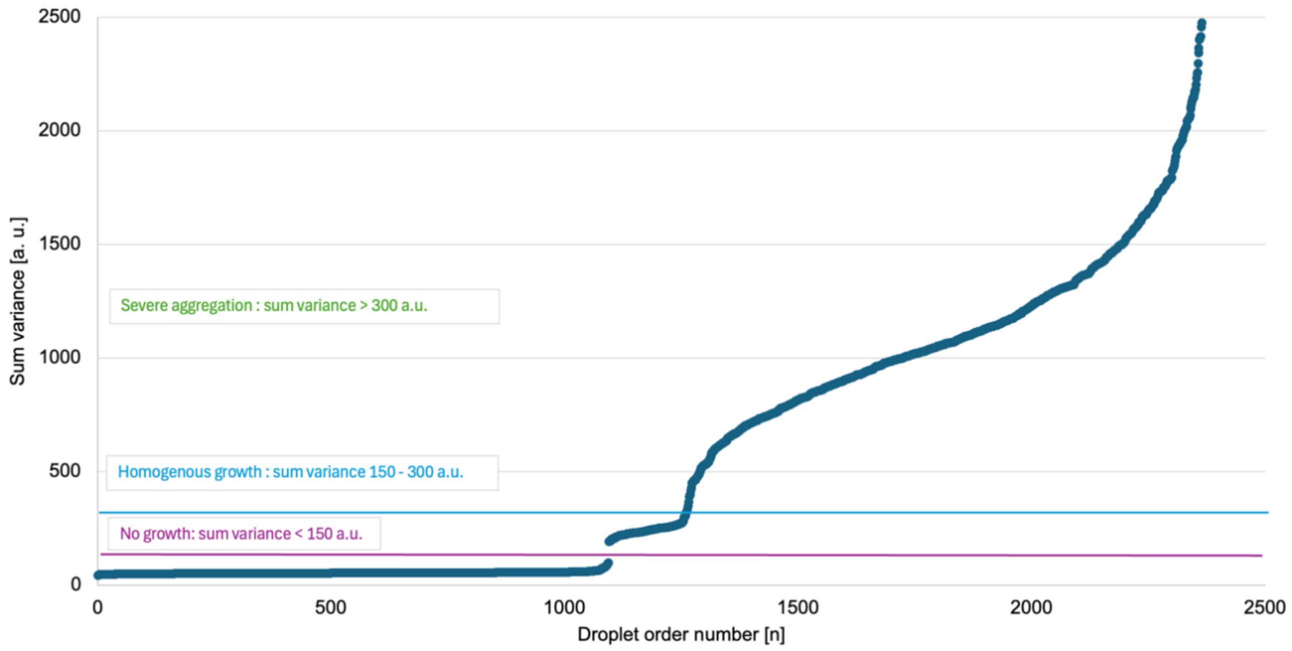


**d** Polydisperse droplet experiment studying bacterial aggregation rates - classification thresholds for cefotaxime dataset

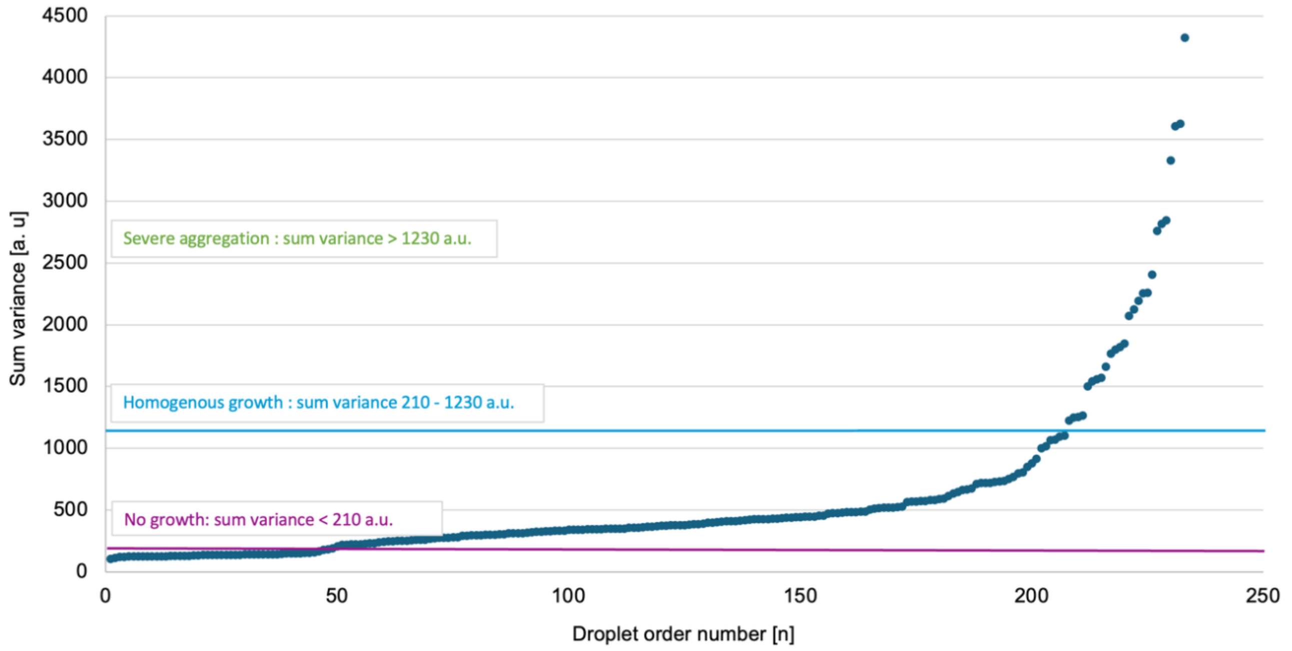

**e** Polydisperse droplet experiment studying bacterial aggregation rates - classification thresholds for chloramphenicol dataset

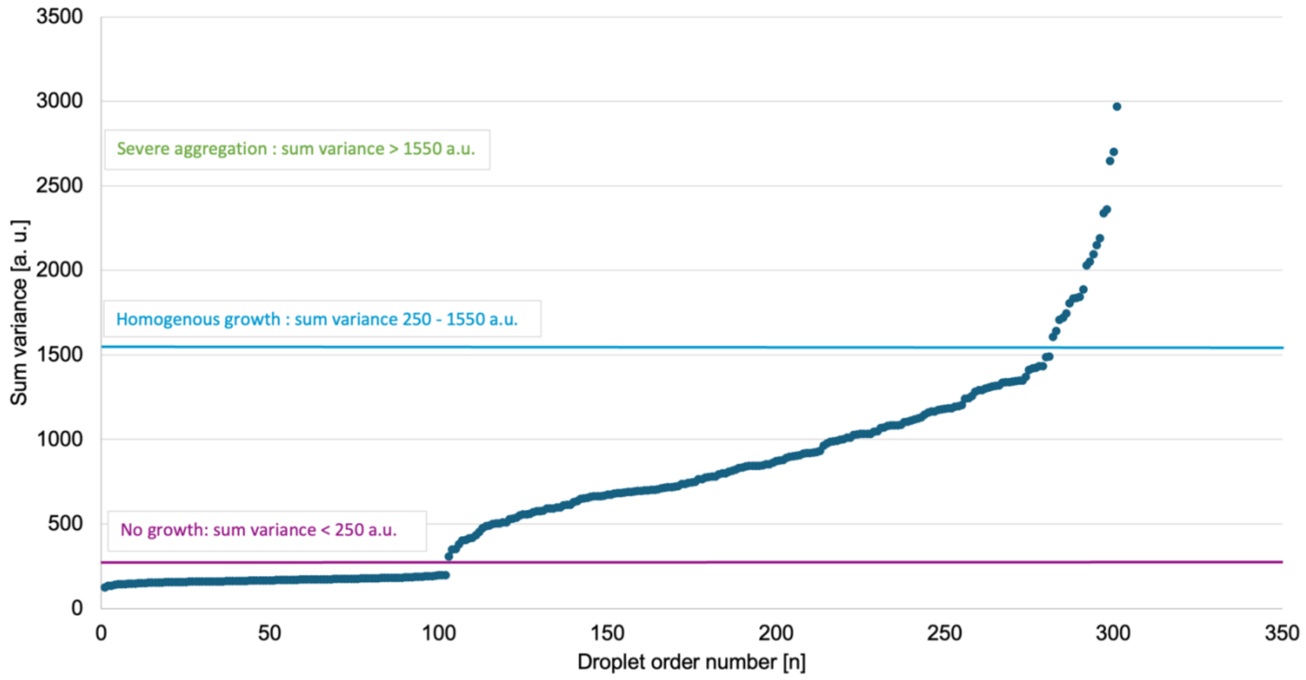


**f** Polydisperse droplet experiment studying bacterial aggregation rates - classification thresholds for ciprofloxacin dataset

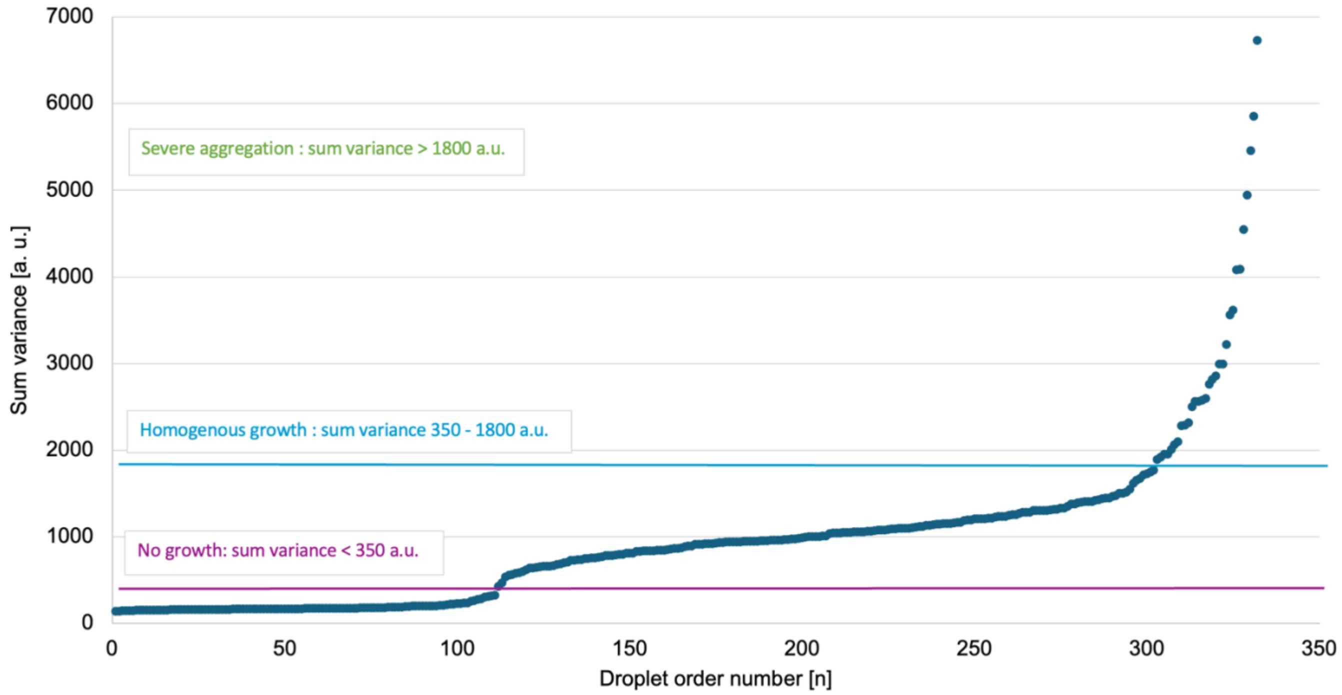

**g** Polydisperse droplet experiment studying bacterial aggregation rates - classification thresholds for doxycycline dataset

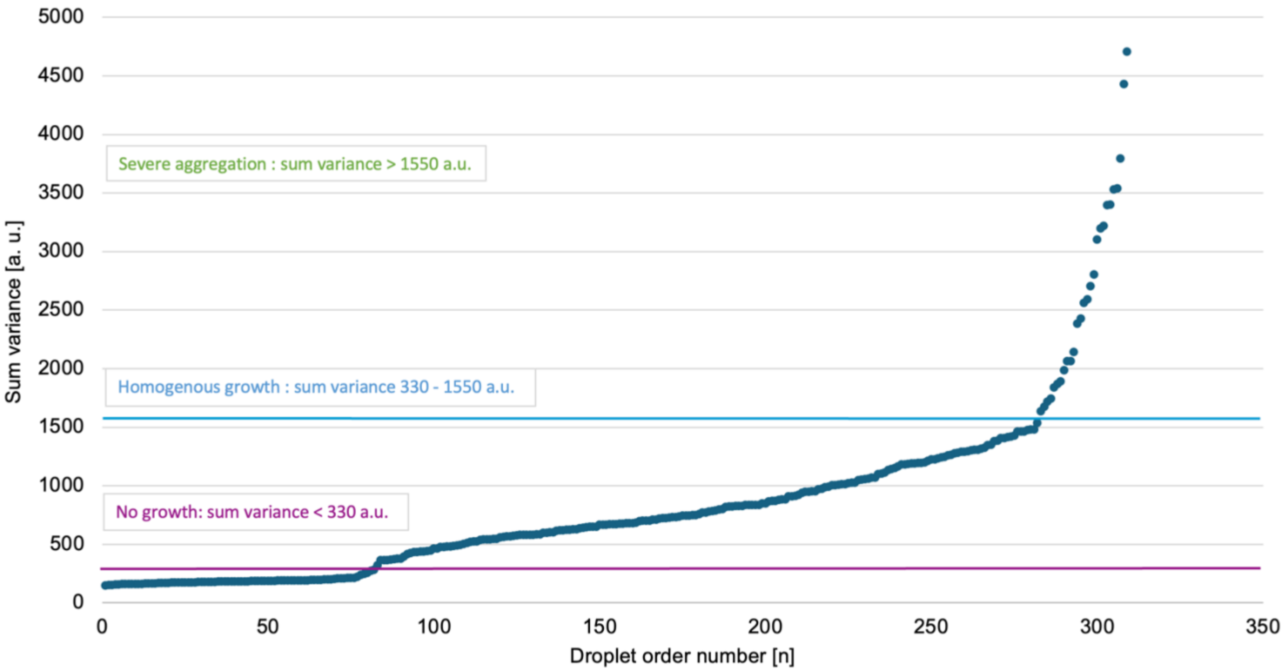


**h** Polydisperse droplet experiment studying bacterial aggregation rates - classification thresholds for trimethoprim dataset

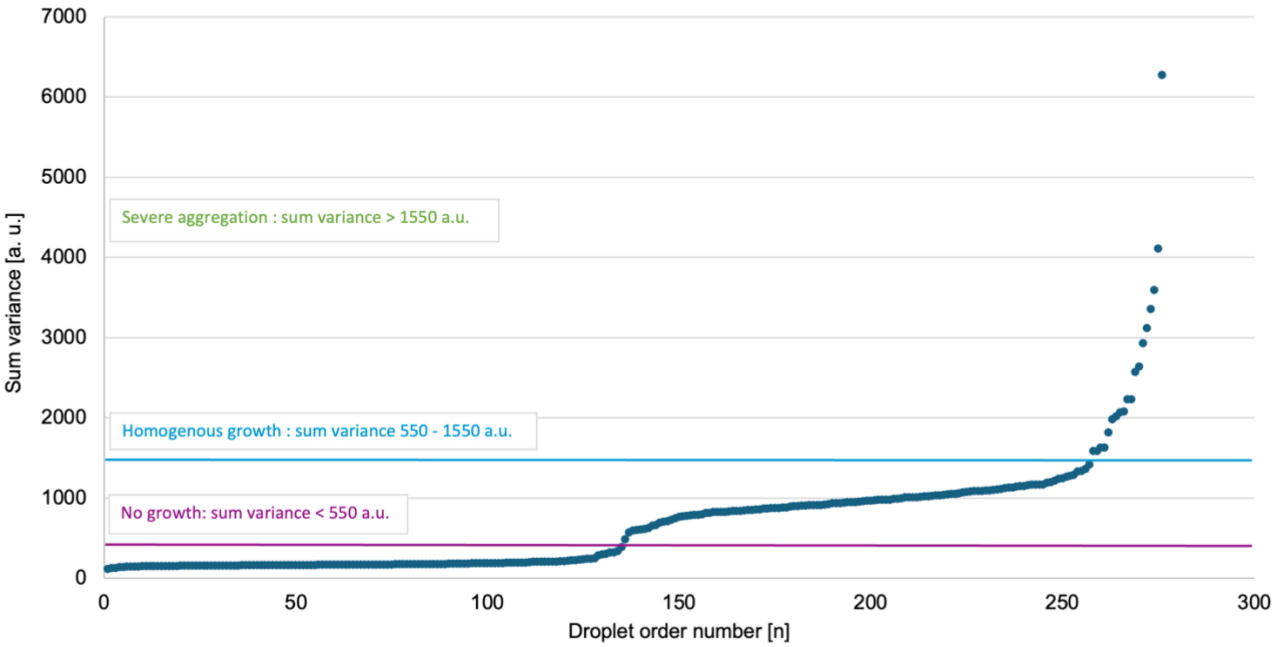

**i** Polydisperse droplet experiment comparing bacterial aggregation rates with and without microplastic presence - classification thresholds for cefotaxime dataset

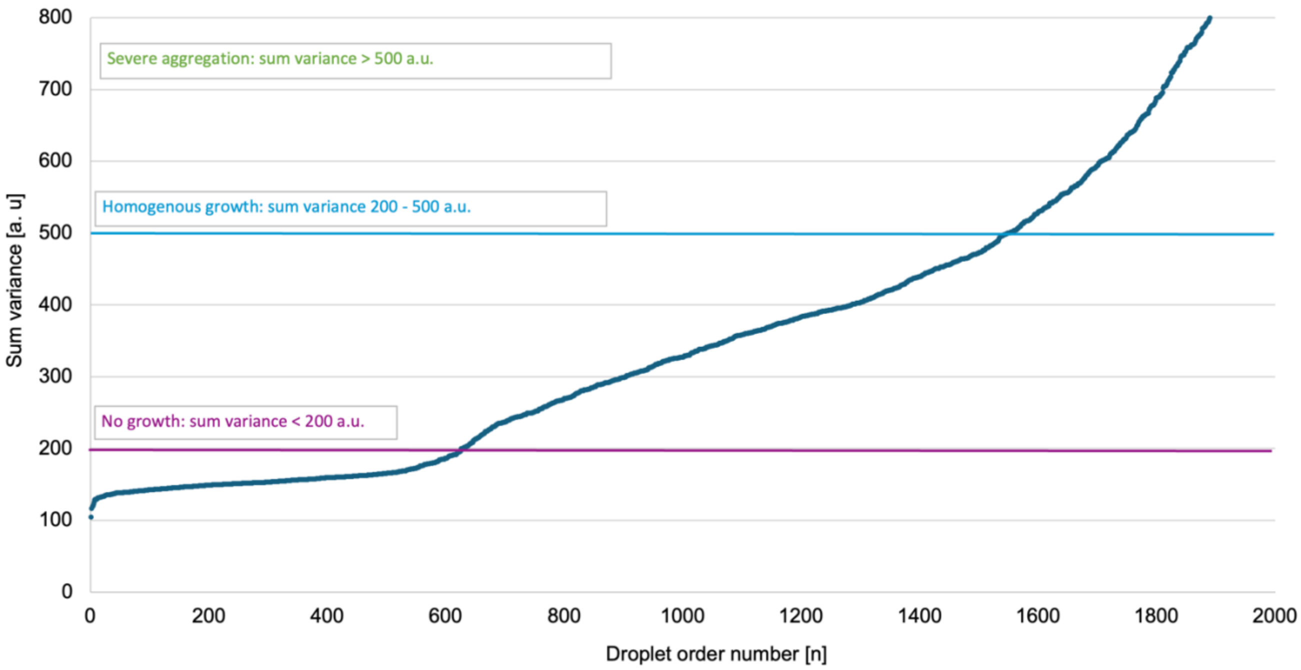


**j** Polydisperse droplet experiment studying bacterial aggregation rates in presence of metals - classification thresholds for $CuSO_4$ dataset

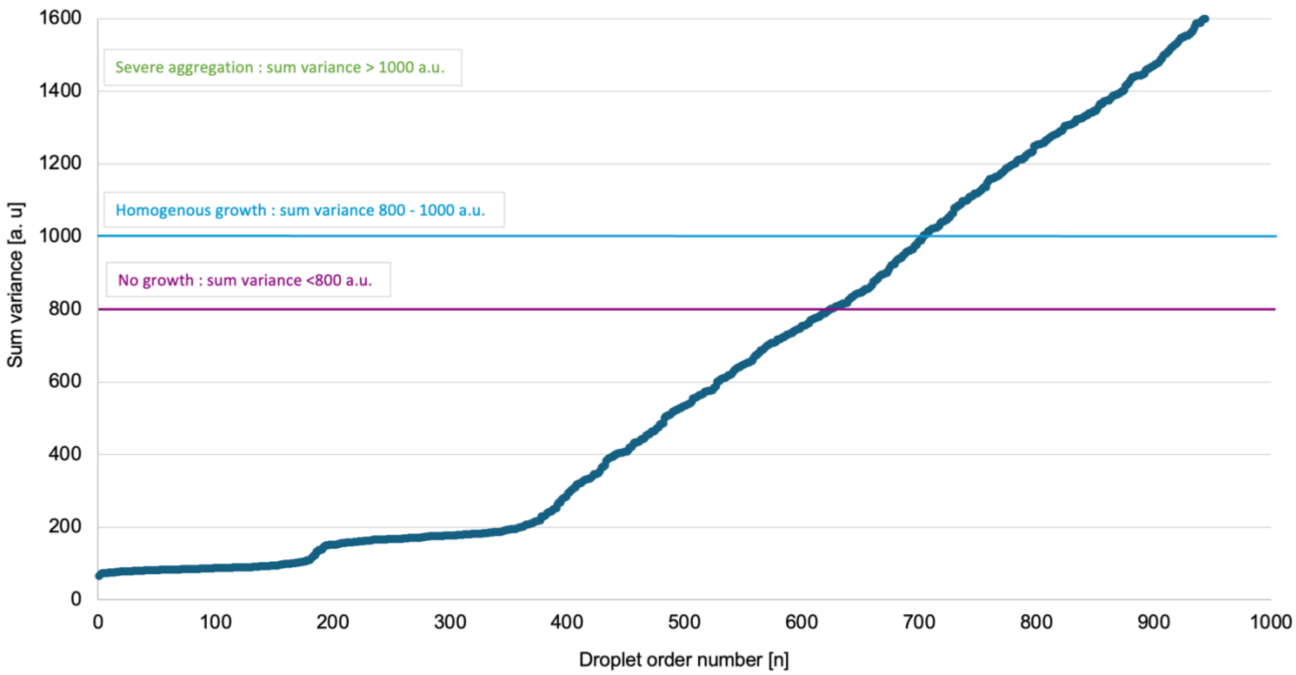

**k** Polydisperse droplet experiment studying bacterial aggregation rates in presence of metals - classification thresholds for LiCl dataset

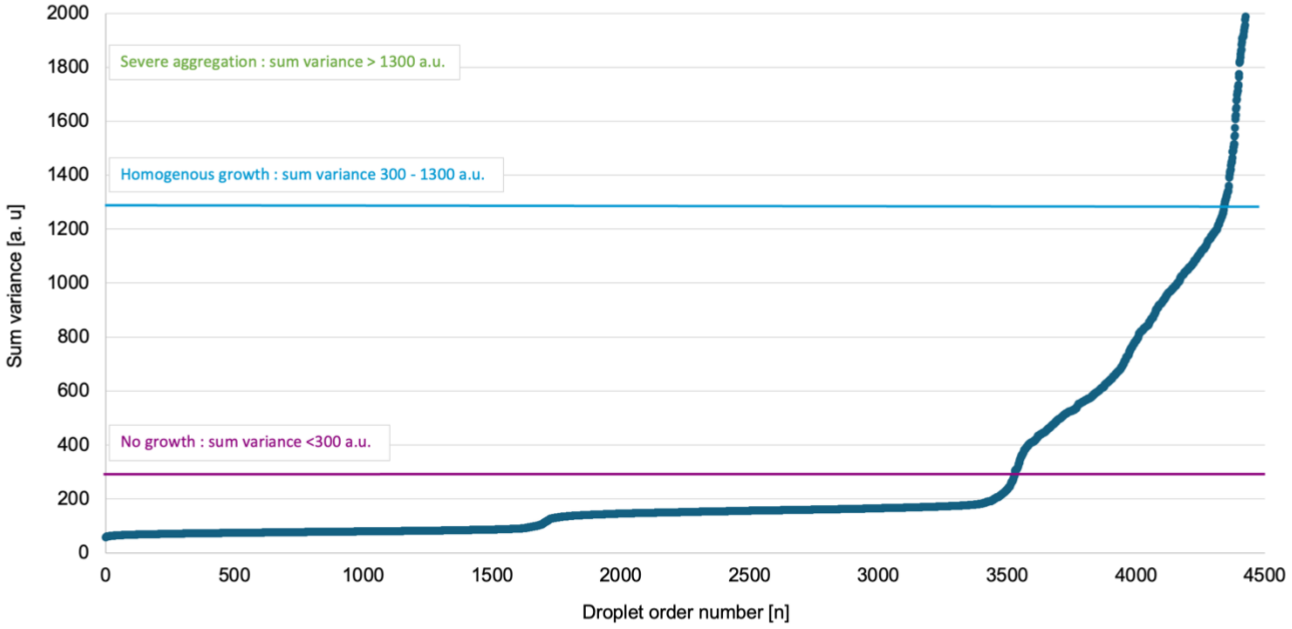


**l** Polydisperse droplet experiments studying bacterial aggregation rates in presence of metals - classification thresholds for $ZnCl_3$

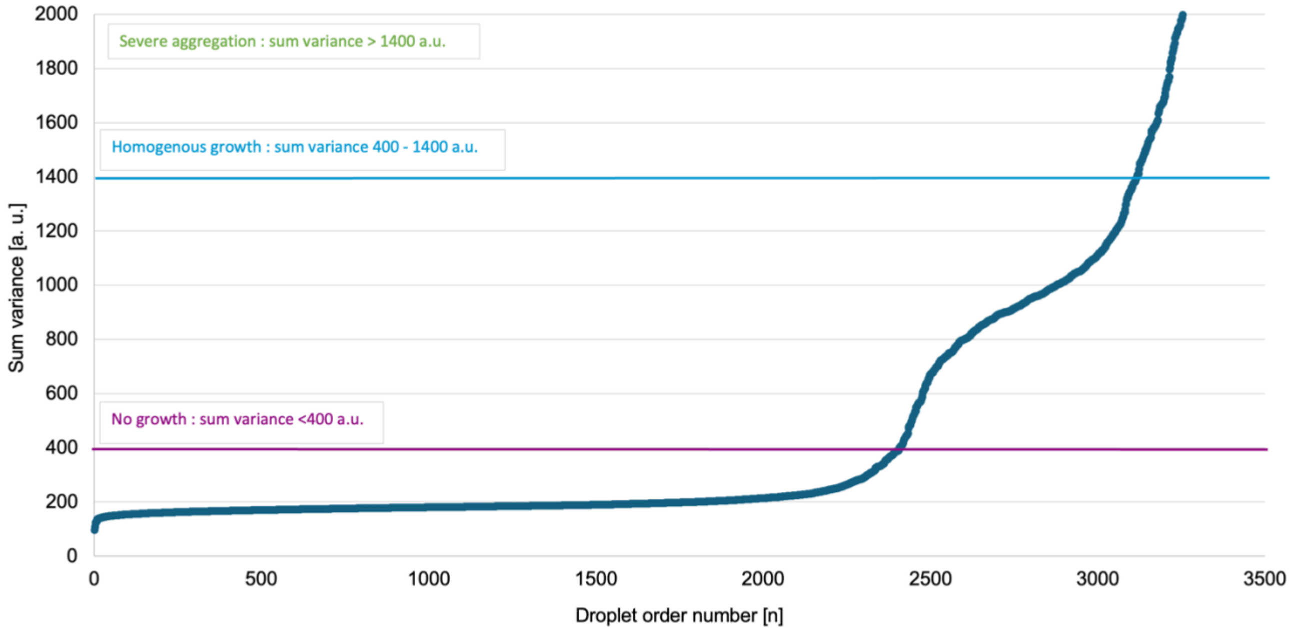

**Table S1.** Performance assessment of reference sets. Accuracy and precision of pipeline was assessed by comparison with a human expert panel for aggregated bacterial growth, homogenous bacterial growth and droplets without bacterial growth.

**a** Droplets with aggregated bacterial growth (reference set)

| | Monodisperse droplets | | Polydisperse droplets | | | | |
|---|---|---|---|---|---|---|---|
| | CTX | KAN | CTX | CHL | CPFX | DOXY | TMP |
| **Accuracy** [%] | **94.8** | **93.5** | **96.1** | **94.0** | **98.5** | **97.1** | **98.6** |
| **Precision** [%] | **82.8** | **78.2** | **100** | **65** | **100** | **85.2** | **89.5** |
| True positive [n] | 53 | 43 | 25 | 13 | 30 | 23 | 17 |
| True negative [n] | 149 | 129 | 199 | 270 | 297 | 277 | 255 |
| False positive [n] | 11 | 12 | 0 | 7 | 0 | 4 | 2 |
| False negative [n] | 0 | 0 | 9 | 11 | 5 | 5 | 2 |
| Total droplets [n] | 213 | 184 | 233 | 301 | 332 | 309 | 276 |

**b** Droplets with homogenous bacterial growth (reference set)

| | Monodisperse droplets | | Polydisperse droplets | | | | |
|---|---|---|---|---|---|---|---|
| | CTX | KAN | CTX | CHL | CPFX | DOXY | TMP |
| **Accuracy** [%] | **94.8** | **93.5** | **92.3** | **93.0** | **96.4** | **94.2** | **97.1** |
| **Precision** [%] | **100** | **100** | **88.6** | **92.2** | **93.7** | **93.0** | **95.0** |
| True positive [n] | 11 | 5 | 140 | 165 | 179 | 185 | 115 |
| True negative [n] | 191 | 167 | 75 | 115 | 141 | 106 | 153 |
| False positive [n] | 0 | 0 | 18 | 14 | 12 | 14 | 6 |
| False negative [n] | 11 | 12 | 0 | 7 | 0 | 4 | 2 |
| Total droplets [n] | 213 | 184 | 233 | 301 | 332 | 309 | 276 |

**c** Droplets without bacterial growth (reference set)

| | Monodisperse droplets | | Polydisperse droplets | | | | |
|---|---|---|---|---|---|---|---|
| | CTX | KAN | CTX | CHL | CPFX | DOXY | TMP |
| **Accuracy** [%] | **100** | **100** | **96.1** | **99.0** | **97.9** | **97.1** | **98.6** |
| **Precision** [%] | **100** | **100** | **100** | **100** | **100** | **100** | **100** |
| True positive [n] | 138 | 124 | 50 | 102 | 111 | 83 | 136 |
| True negative [n] | 75 | 60 | 174 | 196 | 214 | 217 | 136 |
| False positive [n] | 0 | 0 | 0 | 0 | 0 | 0 | 0 |
| False negative [n] | 0 | 0 | 9 | 3 | 7 | 9 | 4 |
| Total droplets [n] | 213 | 184 | 233 | 301 | 332 | 309 | 276 |
